# AI Discovery of Mechanisms of Consciousness, Its Disorders, and Their Treatment

**DOI:** 10.1101/2024.10.16.618720

**Authors:** Daniel Toker, Zhong Sheng Zheng, Jasmine A. Thum, Jing Guang, Jitka Annen, Hiroyuki Miyamoto, Kazuhiro Yamakawa, Paul M. Vespa, Steven Laureys, Caroline Schnakers, Ausaf A. Bari, Andrew Hudson, Nader Pouratian, Martin M. Monti

**Affiliations:** Department of Neurology, University of California, Los Angeles, Los Angeles, CA, USA; Department of Psychology, University of California, Los Angeles, Los Angeles, CA, USA; Research Institute, Casa Colina Hospital and Centers for Healthcare, Pomona, CA, USA; Department of Neurosurgery, Heersink School of Medicine, University of Alabama at Birmingham, Birmingham, AL, USA; Edmond and Lily Safra Center for Brain Sciences, The Hebrew University of Jerusalem, Jerusalem, Israel; Coma Science Group, GIGA-Research, University of Liège, Liège, Belgium; Department of Data Analysis, University of Ghent, Ghent, B9000, Belgium; Laboratory for Neurogenetics, RIKEN Center for Brain Science, Saitama, Japan; PRESTO, Japan Science and Technology Agency, Saitama, Japan; International Research Center for Neurointelligence, University of Tokyo, Tokyo, Japan; Department of Neurodevelopmental Disorder Genetics, Institute of Brain Science, Nagoya City University Graduate School of Medical Science, Nagoya, Japan; Brain Injury Research Center, Department of Neurosurgery, University of California, Los Angeles, Los Angeles, CA, USA; CERVO Brain Research Centre, Laval University & CIUSSS-CN, Québec, Canada; Joint International Research Unit on Neuroplasticity, University of Liège, Liège, Belgium; Anesthesia, Critical Care and Pain Medicine Research, Harvard Medical School, Beth Israel Deaconess Medical Center, Boston, MA, USA; European Foundation of Biomedical Research FERB Onlus, Milano, Italy; Department of Anesthesiology, Veterans Affairs Greater Los Angeles Healthcare System, Los Angeles, CA, USA; Department of Anesthesiology and Perioperative Medicine, University of California, Los Angeles, Los Angeles, CA, USA; Department of Neurological Surgery, UT Southwestern Medical Center, Dallas, TX, USA

**Keywords:** disorders of consciousness, artificial intelligence, neural field theory, deep brain stimulation

## Abstract

Understanding disorders of consciousness (DOC) remains one of the most challenging problems in neuroscience, hindered by the lack of experimental models for probing mechanisms or testing interventions [1]. To address this, we introduce a generative adversarial AI framework [2] that pits deep neural networks—trained to detect consciousness across over 680,000 neuroelectrophysiology samples and validated on 565 patients, healthy volunteers, and animals—against interpretable, machine learning-driven neural field models [3]. This adversarial architecture produces biologically realistic simulations of both conscious and comatose brains that recapitulate empirical neurophysiological features across humans, monkeys, rats, and bats. Without explicit programming, the AI model retrodicts known DOC responses to brain stimulation and generates testable predictions about unconsciousness mechanisms. Two such predictions are validated here: selective disruption of the basal ganglia indirect pathway, supported by diffusion MRI in 51 DOC patients; and increased cortical inhibitory-to-inhibitory synaptic coupling, supported by RNA sequencing from resected brain tissue in six human coma patients and a rat stroke model. The model also identifies high-frequency subthalamic nucleus stimulation as a promising DOC intervention, supported here using electrophysiology data from human patients. This work introduces an AI framework for causal inference and therapeutic discovery in consciousness research and complex systems more broadly.

## INTRODUCTION

Understanding how consciousness arises—and how it collapses in conditions like coma—remains one of the most enduring and difficult challenges in science. Despite major advances in brain imaging, clinicians still lack a mechanistic understanding of consciousness, limiting treatment options for disorders of consciousness (DOC), including coma, the vegetative state, and the minimally conscious state [1]. While therapies such as deep brain stimulation (DBS) have shown promise [4], uncertainty about their mechanisms and optimal targets continues to hinder clinical progress.

A leading hypothesis suggests that consciousness depends on the integrity of a distributed forebrain circuit involving the cortex, thalamus, and basal ganglia [5]. According to this view, injury disrupts inputs to the central thalamus, reducing excitation in cortical and striatal networks, which in turn over-activates the globus pallidus interna (GPi)—a key inhibitory hub that suppresses thalamocortical output. More recent work has also implicated the globus pallidus externa (GPe) as a crucial modulator of consciousness, owing to its influence on arousal-related rhythms and its projections to the cortex and thalamic reticular nucleus (TRN) [6, 7]. Structural imaging studies have further linked GPe atrophy to impaired consciousness [8], yet these circuit-level hypotheses remain difficult to test directly owing to the scarcity of suitable experimental models of DOC.

To address this, we introduce a new artificial intelligence (AI) framework for discovering and validating candidate mechanisms of consciousness. Our approach pits deep neural networks—trained to detect consciousness across more than 680,000 electroencephalography (EEG), electrocorticography (ECoG), and local field potential (LFPs) samples from humans and animals—against machine learning-driven, biophysically grounded dynamical brain models. These models serve as interpretable “generators” in an adversarial loop, and are iteratively tuned via machine learning to produce dynamics that the trained deep neural networks classify as real and conscious. This generative-discriminative architecture enables causal exploration of how specific circuit changes give rise to shifts in consciousness.

We first apply this framework to simulate the healthy conscious brain, generating synthetic neurodynamics that replicate empirical features across humans, monkeys, rats, and bats. We then extend the model to replicate pathological unconsciousness, using networks trained on data from acute coma and chronic DOC patients, and validated on large independent datasets. Without explicit programming, the resulting simulations reproduce known features of DOC and reveal novel mechanisms of impaired awareness. Two predictions—increased synaptic coupling among cortical interneurons, and selective disruption of the basal ganglia indirect pathway—are empirically tested using single-nucleus RNA sequencing and diffusion MRI in patients. Beyond identifying mechanisms, our AI framework enables virtual testing of therapeutic strategies. Without prior knowledge, the model identifies established DBS targets and proposes high-frequency subthalamic stimulation as a promising, previously underexplored intervention. We test this prediction in six human patients receiving subthalamic DBS.

Together, these results demonstrate a general strategy for using AI to uncover mechanistic insight in complex systems. By combining interpretable dynamical models with data-driven optimization, our framework yields testable hypotheses about consciousness and offers a path toward precision therapies for DOC.

## RESULTS

### A generative adversarial AI model of the conscious brain

To identify the circuit-level dynamics that support consciousness, we curated a large-scale, cross-species electrophysiology dataset comprising over 680,000 samples from cortex, thalamus, and basal ganglia in humans, nonhuman primates, and rodents (Supplementary Table 1). Each 10-second sample was labeled with a corresponding behavioral index of consciousness—either a score on the Glasgow Coma Scale (GCS) or Coma Recovery Scale-Revised (CRS-R), or a binary label (conscious vs. comatose). Cortical data (EEG and ECoG) were uniformly preprocessed by resampling to 250 Hz and bandpass filtering from 0.5–45 Hz; subcortical LFPs were sampled at 250 Hz and left unfiltered. Further preprocessing details are provided in the Methods.

To decode consciousness from these signals, we trained deep convolutional neural networks (DCNNs) to output a continuous score from 0 (unconscious) to 1 (fully conscious). Model performance was robust across multiple brain regions, species, and recording modalities. A thalamic DCNN accurately distinguished anesthetic coma from wakefulness in held-out thalamic LFP data from essential tremor (ET) patients (AUC=0.98, 118 validation samples, *n* = 10 patients) and Long-Evans rats (AUC=0.69, 50 validation samples, *n* = 9; Fig. 1B). A GPe-trained DCNN performed similarly in detecting consciousness in validation pallidal LFPs in African green monkeys (AUC=0.96, 3,277 validation samples, *n* = 13) and achieved moderate accuracy in pallidal LFPs from human Parkinson’s disease (PD) patients (AUC=0.84, 24 validation samples, *n* = 12; Fig. 1C).

**Fig. 1:**
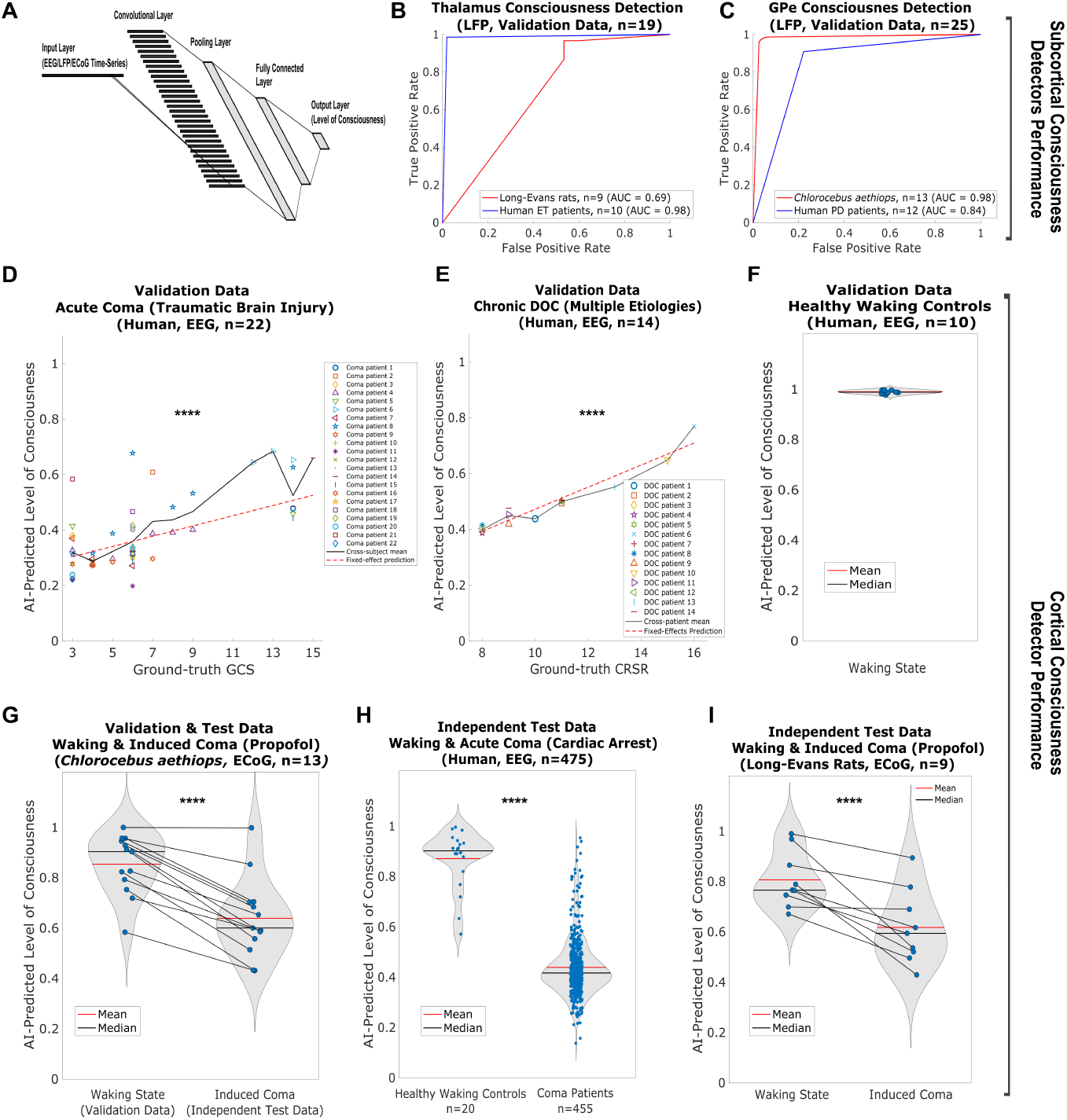
DCNN-based consciousness detection across species and brain regions. (**A**) Architecture of the DCNNs: input layer (10s neuroelectrophysiology time-series), followed by convolutional, pooling, fully connected, and output layers. (**B**) ROC curves for a thalamic DCNN distinguishing wakefulness from anesthetic coma in held-out data from human ET patients (118 samples, *n* = 10, AUC=0.98) and Long-Evans rats (50 samples, *n* = 9, AUC=0.69). (**C**) ROC curves for a GPe DCNN in held-out data from African green monkeys (3,277 samples, *n* = 13, AUC=0.96) and human PD patients (24 samples, *n* = 12, AUC=0.84). (**D**) AI-predicted consciousness vs. GCS in held-out EEG from acute TBI coma patients (150,674 samples, *n* = 22). Black line: cross-subject mean; red dotted: best-fit from two-level linear mixed-effects model. (**E**) AI-predicted consciousness vs. CRS-R in chronic DOC patients (12,266 validation samples, *n* = 14). Lines as in (D). (**F**) Predicted consciousness levels in validation data from healthy awake controls (1,766 samples, *n* = 10). (**G**) Predictions across wake (1,914 validation samples) and propofol coma (9,570 test samples) in African green monkeys (*n* = 13). (**H**) Predicted consciousness level in independent EEG data from healthy controls (3,674 test samples, *n* = 20) and post–cardiac arrest coma patients (1,180,257 samples, *n* = 455). (**I**) Predicted consciousnes level in waking vs. propofol coma ECoG from Long-Evans rats (250 samples, *n* = 9). Each point in D–I shows the subject-level mean across all 10s samples and channels. Horizontal lines (G, I) link conditions within subjects. Violin plots show distributions; red and black bars mark group mean and median. Statistical comparisons used two-level linear mixed-effects models. D–E used continuous GCS/CRS-R scores; others used binary labels. ****: *p <* 0.0001 (Bonferroni-corrected).

For cortical signals, a DCNN was trained on 602,696 EEG samples from acute TBI coma patients (*n* = 22), 49,062 samples from chronic DOC patients (*n* = 14), 7,066 samples from healthy controls (*n* = 10), and smaller ECoG datasets from ET and PD patients and African green monkeys (see Supplementary Table 1). In held-out EEG validation data, predicted consciousness levels scaled significantly with ground-truth GCS scores in TBI patients (*p <* 0.0001; Fig. 1D) and CRS-R scores in chronic DOC patients (*p <* 0.0001; Fig. 1E). In healthy controls, predicted consciousness was near-maximal (1,766 validation samples, *n* = 10; Fig. 1F). The cortical DCNN also generalized robustly to fully independent datasets: predicted consciousness was significantly reduced in induced propofol coma in African green monkeys (*n* = 13, *p <* 0.0001; Fig. 1G), in post–cardiac arrest coma patients (*n* = 455) compared to healthy controls (*n* = 20; *p <* 0.0001; Fig. 1H), and in Long-Evans rats under propofol (*n* = 9; *p <* 0.0001; Fig. 1I). All statistical comparisons were conducted using two-level linear mixed-effects models to account for nested sampling, with Bonferroni correction for multiple comparisons. A subsequent analysis of the learned filters in the cortical DCNN showed that it prioritizes temporal features in the delta to alpha range (0.5–15 Hz), with a smaller cluster of filters tuned to beta activity (30–35 Hz), suggesting that the network distinguishes conscious from unconscious brain states based on specific temporal patterns within these frequency bands (Supplementary Fig. 1).

To create a generative AI model of the conscious brain, we used these trained DCNNs as discriminators in an adversarial optimization loop. Instead of using a deep neural network as the generator, we employed a biologically grounded neural field model, whose parameters were updated by a genetic algorithm. This choice was motivated by the interpretability of the model: unlike standard neural network generators, the mean-field framework allows for direct mechanistic insights and the generation of testable hypotheses about consciousness. Genetic optimization was chosen for its ability to navigate high-dimensional, non-convex parameter spaces with unknown gradients. Model connectivity was based on known anatomy [9, 10], but connection strengths and biophysical parameters were tuned through optimization. Additional DCNNs trained to detect seizures and distinguish real neural from synthetic signals, along with empirical constraints on firing rates, phase-amplitude coupling patterns, cross-frequency information transfer, and Lyapunov exponents, further guided optimization (see Methods). Once a model output was reached that was classified as conscious, real, and non-seizing by the DCNNs, new discriminator networks were trained to distinguish real from simulated conscious brains. The model was then further refined through genetic optimization until these new DCNNs failed to distinguish simulations of the conscious brain from the real conscious brain (Fig. 2A). Final parameters are listed in Tables S2–S4.

**Fig. 2:**
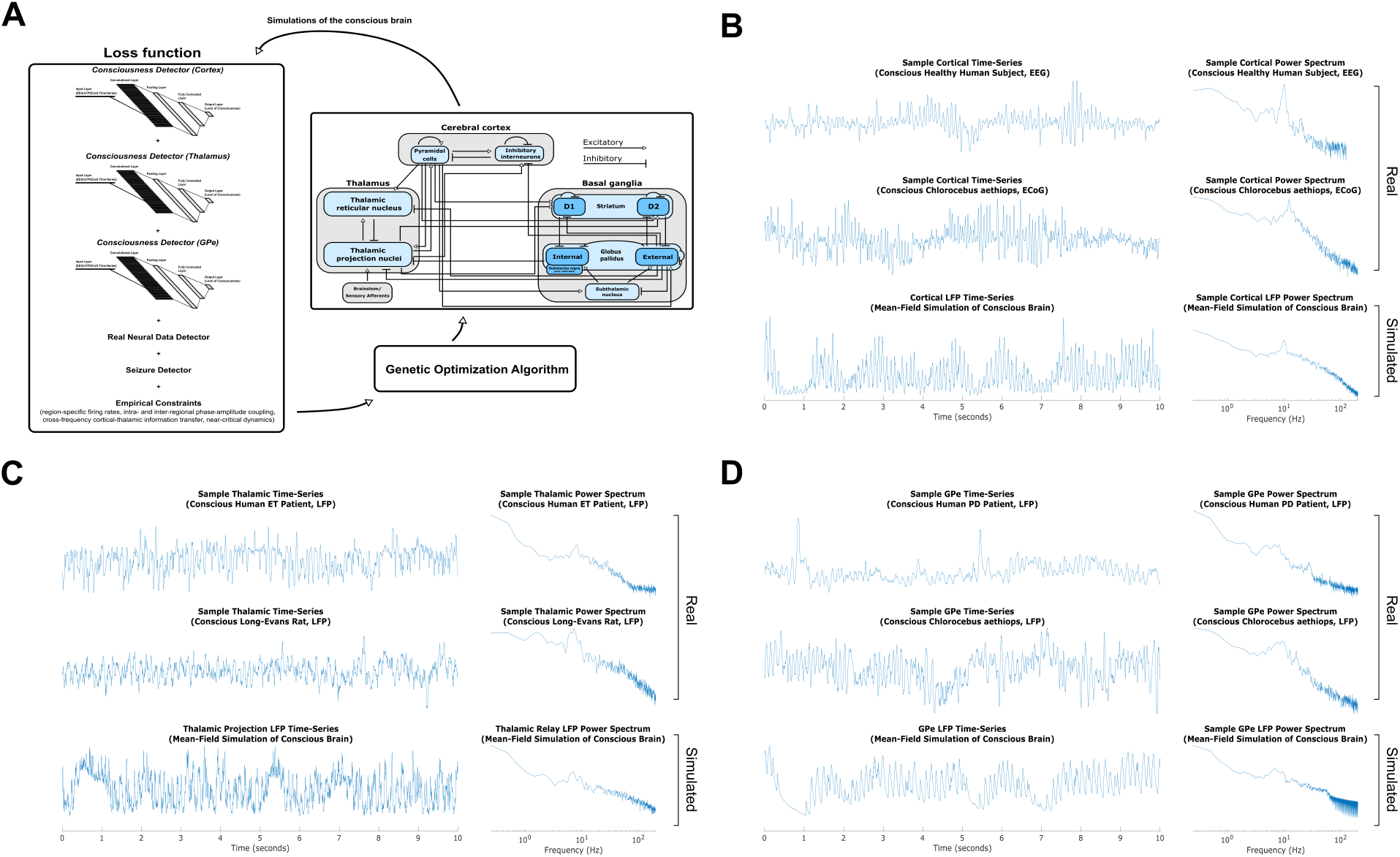
A generative AI model of the conscious thalamo-cortical-basal ganglia system. **(A)** Schematic illustrating the adversarial-like process of using trained deep neural networks and genetic optimization to tune the parameters of a neural field model based on known anatomical connectivity. The loss function incorporated outputs from networks trained to classify real versus synthetic data, detect consciousness, and identify seizures, as well as empirical constraints. **(B-D)** Simulated LFPs (left panels) and power spectra (right panels) from the cortex (B), thalamic projection nuclei (C), and GPe (D), compared to sample real LFPs and their corresponding power spectra from multiple mammalian species.

**Fig. 3:**
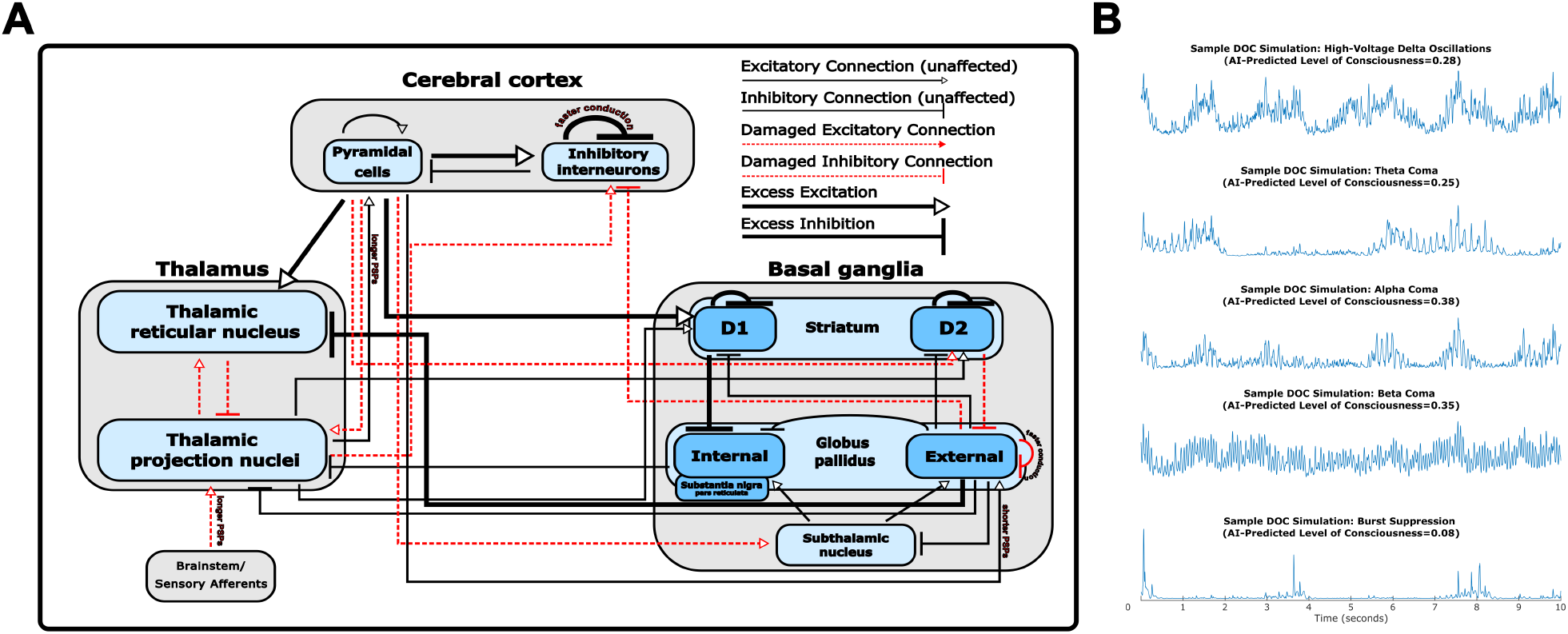
An AI-driven mesocircuit model of disorders of consciousness. **(A)** A visualization of mesocircuit-level changes predicted by our AI model to drive unconsciousness in DOC, corresponding to the changes described in Table 1. **(B)** Sample cortical LFPs from our generative AI models of DOC, matching known electrophysiology phenotypes in these disorders [12]. All LFPs are plotted on the same scale.

LFPs and corresponding spectra from our model of the conscious brain closely matched recordings from rats, monkeys, and humans during wakefulness (Fig.2, Supplementary Fig. 2). Region-specific firing rates were within empirical bounds (Supplementary Table 5). The model also reproduced key neurophysiological signatures of consciousness: theta–gamma phase-amplitude coupling within and between cortex, thalamus, and GPe (Supplementary Fig. 3A–B); bidirectional cross-frequency information transfer between cortex and thalamus (Supplementary Fig. 3C–D) [9]; and cortical dynamics near edge-of-chaos criticality (mean Lyapunov exponent = 0.69; Supplementary Fig. 3E) [9, 11]. Although striatal signals were not used in training, simulated striatal LFPs generalized well to dorsal striatum data from rats and bats, replicating theta–gamma coupling and spectral features (Supplementary Fig. 4). Finally, without explicit programming, features which differentiate real striatal vs. pallidal vs. thalamic LFPs across humans, monkeys, rats, and bats during conscious states were recapitulated in the model (Supplementary Table 6). These results affirm the biological fidelity of our generative AI model of the conscious brain.

### An adversarial AI approach identifies circuit-level drivers of DOC

To probe the mechanisms underlying DOC, we reprogrammed our generative AI model to optimize for unconsciousness, flipping its objective from simulating waking brain activity to replicating the dynamics of comatose states. Starting from our optimized model of the conscious brain, we applied genetic optimization with randomized initializations to generate 280 distinct DOC models. Each was constrained to produce a graded reduction in AI-predicted consciousness with further deviation from the “conscious brain” model (as measured by our cortical DOC-trained classifier), while being classified by our trained DCNNs as both real and non-seizing. The resulting in silico DOC cohort exhibited a rich diversity of electrophysiological phenotypes—including generalized slowing and burst suppression—that mirror clinical observations in coma and vegetative states [12].

To identify circuit parameters most predictive of consciousness level, we modeled six degrees of “severity” for each of our 280 in silico DOC models (yielding 1,680 simulations total), and regressed all AI-estimated consciousness scores against model features ranging from synaptic coupling to conduction delays and synaptic kinetics. All variables were rank-normalized via inverse Gaussian transform, and significance was determined using empirical p-values from 1,000 permutations, with FDR correction applied separately to classes of model features. The results, summarized in Table 1, reproduce key elements of the mesocircuit hypothesis [5, 13], including weakened cortical drive to thalamus and striatum and disruption of the striatopallidal pathway. For a more detailed discussion of the model’s alignment with known features of DOC and severe brain injury, see Discussion. In addition to successful retrodiction, our AI model also yields novel predicted drivers of unconsciousness: notably, increased inhibitory-to-inhibitory synaptic coupling within cortex, and selective degradation of projections from D2-expressing cells in the striatum to the GPe, with moderate strengthening of pathways from D1-expressing striatal cells to the GPi. Additional novel predicted contributors to unconsciousness included prolonged corticothalamic synaptic integration, reduced subcortical input to cortical interneurons, and faster conduction among inhibitory neurons.

**Table 1:** Parameters, regional firing rates, and dynamical features within the generative AI DOC model that are significantly associated with DCNN–predicted level of consciousness. A rank-based inverse Gaussian transformation was applied to all features and to the AI-predicted consciousness levels prior to ridge-penalized linear regression. Empirical p-values were calculated using 1,000 permutations of the AI-predicted consciousness levels, and FDR correction was applied separately within each feature category. Each coefficient reflects the standardized association between a feature and predicted level of consciousness. “Coupling” denotes axonal connection strength. “Striatum (D1-expressing)” and “Striatum (D2-expressing)” refer to specific populations of striatal medium spiny neurons expressing D1 or D2 dopamine receptors, respectively. Novel predicted correlates of consciousness in DOC, which we here empirically test, are bolded.

| **In Silico Axonal Connections Associated with AI-Predicted Level of Consciousness** | **Coefficient**<br>(Ridge regression) | **p-value**<br>(*Permutation test*)<br>(*FDR-corrected*) |
| --- | --- | --- |
| **Inhibitory Cortical to Inhibitory Cortical Coupling** | -0.270 | <0.0001 |
| **Striatum (D2-expressing) to GPe Coupling** | +0.160 | <0.0001 |
| TRN to Thalamic Projection Coupling | +0.123 | <0.0001 |
| Brainstem/Sensory Afferent to Thalamic Projection Coupling | +0.122 | <0.0001 |
| Excitatory Cortex to Striatum (D1-expressing) Coupling | -0.119 | 0.0058 |
| Excitatory to Inhibitory Cortical Coupling | -0.109 | 0.0058 |
| **Striatum (D1-expressing) to GPi Coupling** | -0.105 | 0.0180 |
| Excitatory Cortex to Thalamic Projection Coupling | +0.093 | 0.0100 |
| Striatum (D2-expressing) to Striatum (D2-expressing) Coupling | -0.089 | 0.020 |
| Striatum (D1-expressing) to Striatum (D1-expressing) Coupling | -0.088 | 0.031 |
| GPe to GPe Coupling | +0.085 | 0.0058 |
| Thalamic Projection to Inhibitory Cortex Coupling | +0.084 | 0.0280 |
| Excitatory Cortex to TRN Coupling | -0.084 | 0.0380 |
| GPe to TRN Coupling | -0.084 | 0.0260 |
| GPe to Inhibitory Cortex Coupling | +0.083 | 0.0360 |
| Excitatory Cortex to STN Coupling | +0.082 | 0.0350 |
| Excitatory Cortex to Striatum (D2-expressing) Coupling | +0.079 | 0.0360 |
| Thalamic Projection to TRN Coupling | +0.071 | 0.0480 |
| **In Silico Biophysical Features Associated with AI-Predicted Level of Consciousness** |  |  |
| Inhibitory Cortical to Inhibitory Cortical Propagation Delay | +0.466 | <0.0001 |
| Thalamic Projection to Excitatory Cortical PSP Duration | -0.283 | <0.0001 |
| Excitatory Cortical to GPe PSP Duration | +0.230 | <0.0001 |
| Brainstem/Sensory Afferent to Thalamic Projection PSP Duration | -0.220 | <0.0001 |
| GPe to GPe Propagation Delay | +0.185 | <0.0001 |
| **In Silico Regional Firing Rates Associated with AI-Predicted Level of Consciousness** |  |  |
| Excitatory Cortical Firing Rate | +0.461 | <0.0001 |
| GPe Firing Rate | -0.307 | <0.0001 |
| Inhibitory Cortical Firing Rate | -0.239 | <0.0001 |
| TRN Firing Rate | -0.211 | <0.0001 |
| Thalamic Projection Firing Rate | +0.155 | 0.0016 |
| **In Silico Dynamical and Information-Theoretic Features Associated with AI-Predicted Level of Consciousness** |  |  |
| Permutation Entropy ($\tau=32$) | +0.241 | <0.0001 |
| Distance from Edge-of-Chaos Criticality | -0.241 | <0.0001 |
| Multiscale Sample Entropy | +0.181 | <0.0001 |
| Alpha power | +0.173 | <0.0001 |
| Thalamocortical Cross-Frequency Information Transfer | +0.162 | <0.0001 |
| Lempel-Ziv Complexity | +0.116 | 0.027 |
| Permutation Entropy ($\tau=16$) | +0.078 | 0.013 |

Firing rate patterns also echoed known hallmarks of coma: reduced activity in cortical and thalamic projection neurons tracked lower predicted consciousness, while elevated firing in pallidal and TRN populations tracked lower scores (Table 1) (see Discussion for comparison to empirical results). Moreover, replicating empirical data in DOC [14], alpha power was a strong positive predictor (*β* = 0.432, *p <* 0.001) of conscious level, while delta power was negatively associated (*β* = −0.067, *p* = 0.005), though only alpha survived in our multivariate regression (Table 1). Among complexity measures, which consistently positively correlate with level of consciousness empirically [11, 15], permutation entropy in the *β*-range (*τ* =32) and proximity to edge-of-chaos criticality [9, 11] were the strongest independent predictors of consciousness in our generative models. Low-to-high frequency directed information transfer from thalamus to cortex also tracked predicted consciousness levels (Table 1), consistent with prior empirical results [9].

### RNAseq validation of AI-predicted up-regulation of cortical inhibitory to inhibitory synaptic coupling in DOC

One of the top predicted drivers of pathological unconsciousness in our generative AI model was increased inhibitory-to-inhibitory synaptic coupling in cortex (Table 1). To empirically test this prediction, we analyzed single-nucleus RNA sequencing (snR-NAseq) data from resected cortex of patients in acute coma (Glasgow Coma Scale *≤*8) following traumatic brain injury, focusing specifically on parvalbumin-positive (PV^+^) interneurons—the same fast-spiking subclass modeled in our generative framework (mean firing rate *∼*18 Hz; Supplementary Table 5). Given that up-regulation of *VGF* and *SCG2* drives cortical PV^+^ *→*PV^+^ synaptogenesis [16], we tested their expression using a one-tailed GLMM with donor as a random intercept. Both genes were significantly up-regulated in cortical PV^+^ cells from coma patients: *VGF* (Δlog-odds = 1.70 *±* 0.52, *t* = 3.29, *df* = 191, *p* = 0.0012) and *SCG2* (Δlog-odds = 1.24 *±* 0.50, *t* = 2.49, *df* = 191, *p* = 0.0137) (Fig. 4B–C).

**Fig. 4:**
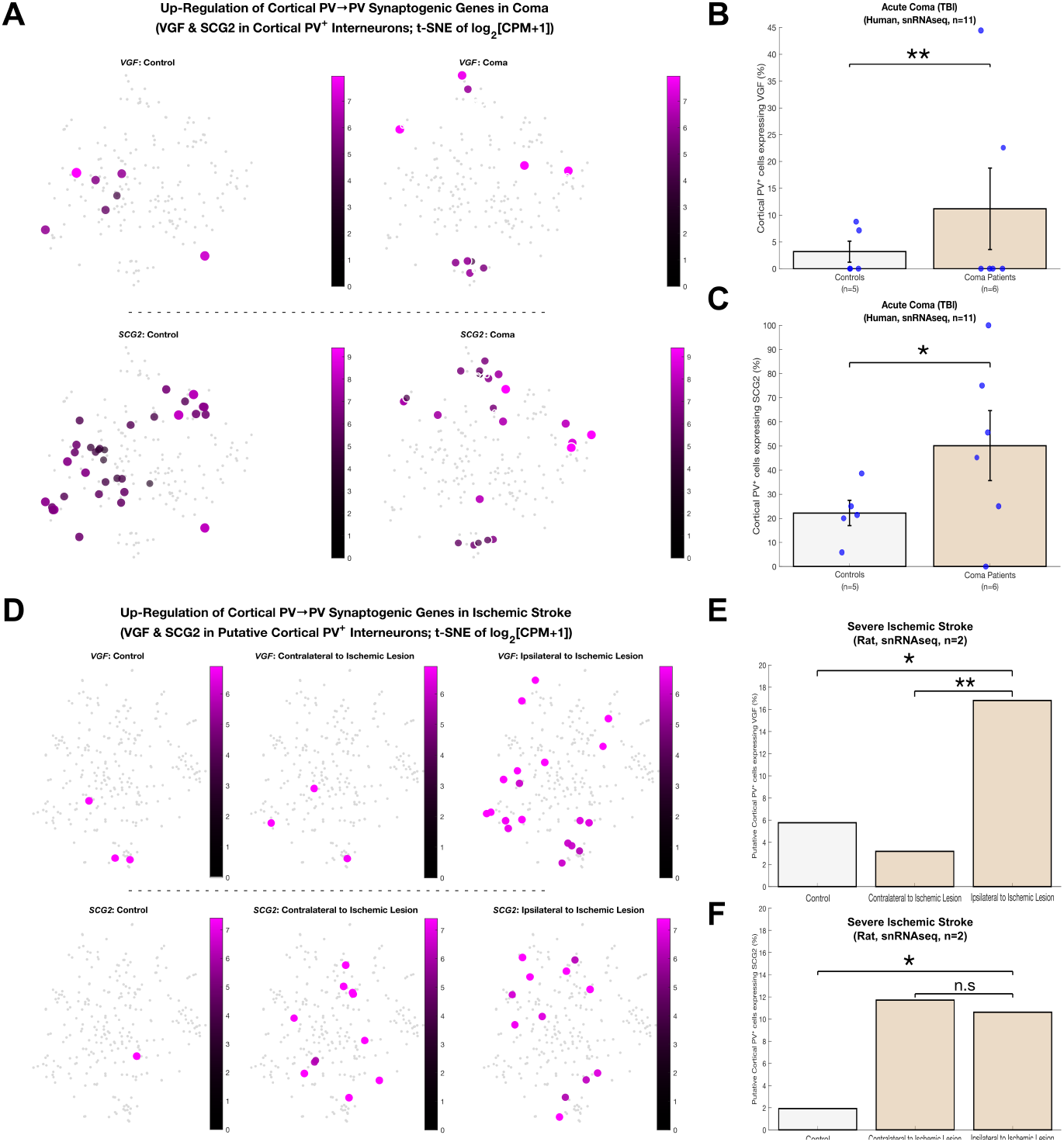
Up-regulation of cortical PV *→*PV synaptogenic genes, an AI-predicted driver of pathological unconsciousness, in acute traumatic coma and severe ischemic stroke. **(A)** Two-dimensional t-distributed stochastic neighbor embedding (t-SNE) of log_2_(counts per million + 1) in human cortical PV^+^ interneurons, showing *VGF* (top) and *SCG2* (bottom) expression in non-neurological controls (left) and acute traumatic coma patients (right). Dot size, opacity, and color encode per-cell expression; background PV^+^ cells are shown in light gray. **(B–C)** Donor-level bar plots of the percentage of PV^+^ cells expressing *VGF* (B) and *SCG2* (C) in controls (n=5, light gray) versus coma patients (n=6, orange). Bars: mean±SEM; overlaid blue dots: individual donors; \*\**p <* 0.01, \**p <* 0.05 (one-tailed GLMM with donor as random effect). **(D)** t-SNE of log_2_(CPM+1) in rat putative cortical PV^+^ interneurons, showing *VGF* (top) and *SCG2* (bottom) expression in sham-operated control (left, light gray), contralateral hemisphere to severe middle cerebral artery occlusion (sMCAO) ischemic stroke (center), and the hemisphere ipsilteral to severe ischemic stroke (right). Visual conventions as in (A). **(E–F)** Sample-level bar plots of the percentage of putative cortical PV^+^ cells expressing *VGF* (E) and *SCG2* (F) in sham control (light gray), hemisphere contralateral to severe ischemic stroke (orange), and hemisphere ipsilateral to severe ischemic stroke (orange) (n=1 sham, n=1 paired stroke). Bars: mean; \**p <* 0.05, \*\**p <* 0.01, n.s. not significant (one-tailed GLMM at the cell level).

To assess generalizability, we analyzed snRNAseq from a rat 48 h following severe ischemic (middle cerebral artery) stroke, a leading cause of coma in humans [17]. Cortical PV^+^ cells were identified by co-expression of *PVALB* and *SATB1* and excluding subcortical markers *PTHLH* and *C1QL1*. In cortex ipsiliteral to the severe ischemic insult, *VGF* was significantly up-regulated relative to control (Δlog-odds = 1.194 *±* 0.646, *t* = 1.850, *df* = 256, *p* = 0.0328) and contralateral hemisphere (Δlog-odds = 1.813, *p* = 0.00243; Fig. 4E). *SCG2* was also up-regulated vs. control (Δlog-odds = 1.802 *±* 1.055, *t* = 1.708, *df* = 256, *p* = 0.0444), but not vs. contralateral hemisphere (Δlog-odds = −0.109, *p* = 0.597; Fig. 4F). As contrasts addressed distinct hypotheses, no correction for multiple comparisons was applied. Other inhibitory synapse–related genes showed mixed patterns across these severe brain injury etiologies (Supplementary Fig. 5). The consistent up-regulation of *VGF* and *SCG2* —which are sufficient to drive cortical PV^+^ *→* PV^+^ synapse formation [16]—across both datasets supports the AI-predicted increase in cortical PV^+^ *→*PV^+^ synaptic coupling in DOC.

### Diffusion tensor imaging validation of AI-predicted disruption to basal ganglia indirect pathway in DOC

To test a second key novel prediction from our generative AI model—that selective disrupted connectivity along the indirect basal ganglia pathway, specifically from D2-expressing striatal neurons to the GPe, partly drives impaired awareness in DOC (Table 1)—we analyzed diffusion tensor imaging (DTI) scans from 51 DOC patients. We quantified structural connectivity between the striatum and each segment of the globus pallidus (GPe and GPi) using probabilistic tractography, calculating the number of streamlines from the striatum that reached each pallidal target.

Patients were grouped by CRS-R diagnosis into vegetative state (VS, *n* = 25) and minimally conscious/emerging MCS (MCS^+^/eMCS, *n* = 29). As predicted, left striatum *→*GPe streamline counts were significantly lower in VS (one-tailed Wilcoxon, *p* = 0.0294, FDR-corrected; Cliff’s delta = −0.3605; Fig. 5A), with a similar trend on the right (*p* = 0.0728; Cliff’s delta = −0.2414; Fig. 5B). A rank-based ANCOVA controlling for age, gender, days post-injury, and etiology supported reduced left striatum *→*GPe streamlines in VS (*β* = −7.82, *p* = 0.0714), despite the significant variance explained by etiology (Supplementary Table 8). After geometric normalization for striatal and pallidal volume, left striatum *→*GPe structural connectivity remained lower in VS (*p* = 0.0540; Cliff’s delta = −0.2665). Striatum*→*GPi counts did not differ by group (Supplementary Fig. 6A–B), but left striatum*→* GPe counts correlated with CRS-R (*ρ* = 0.263, *p* = 0.034, Supplementary Fig. 6C). Fractional anisotropy was unchanged along GPe tracts (Supplementary Fig. 6G–H), but elevated in VS along left striatum*→* GPi (Supplementary Fig. 6I), consistent with model predictions of striatum-GPi pathway strengthening in pathological unconsciousness.

**Fig. 5:**
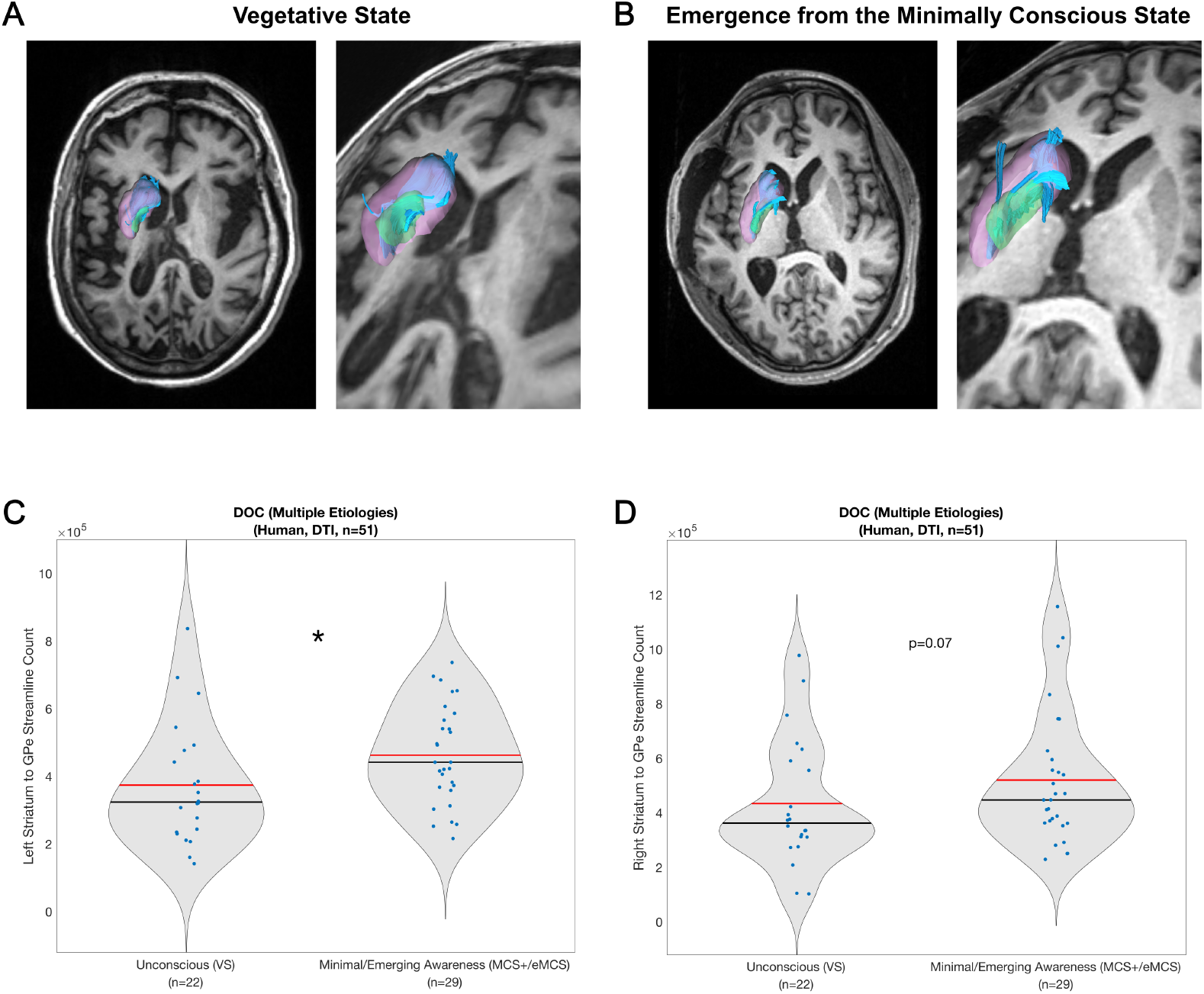
Structural connectivity between the striatum and GPe is reduced in vegetative state patients, matching AI predictions. **(A–B)** Example deterministic tractography reconstructions from diffusion MRI data in two individual DOC patients. Tractography was performed in each patient’s native diffusion space and overlaid on their individual T1-weighted anatomical scan. The striatum (pink) and GPe (green) were defined using subject-specific ROIs. Blue streamlines depict fibers projecting from the left striatum to the left GPe. For each patient, the left image shows an inferior view of the whole brain, while the right image shows a zoomed-in, oblique view focused on the basal ganglia. In both views, the left hemisphere is on the left side of the image. **(A)** Female patient in a vegetative state (VS) of unspecified non-traumatic etiology, 4.61 years post-injury. **(B)** Male patient in emergence from the minimally conscious state (eMCS) following traumatic brain injury, 1 month post-injury. **(C–D)** Group-level differences in striatum-to-GPe structural connectivity in DOC patients (*n* = 51), grouped by diagnosis: vegetative state (VS, *n* = 22) vs. minimally conscious or emerging MCS (MCS+/eMCS, *n* = 29). **(C)** Left hemisphere streamline counts were significantly lower in VS patients compared to MCS+/eMCS (*p* = 0.0294, one-tailed Wilcoxon rank-sum test, FDR-corrected). **(D)** A similar trend was observed in the right hemisphere (*p* = 0.0728, FDR-corrected). Each blue dot represents an individual patient; red and black bars denote the group mean and median, respectively.

### The STN is a promising target for restoring consciousness

Building on our AI model of DOC, we systematically explored the effects of deep brain stimulation (DBS) across a range of subcortical targets and stimulation frequencies (2–130 Hz). To simulate DBS, we incorporated an additional population into the neural field model that generated square-wave pulse trains at defined frequencies. This DBS population lacked dendritic inputs and delivered fixed-strength excitatory output to a specified subcortical target.

For each stimulation condition—defined by a unique pairing of subcortical target and stimulation frequency—we simulated LFPs from the cortex, thalamic projection nuclei, and GPe. These simulated signals were evaluated by our trained cortical, thalamic, and pallidal deep neural network–based consciousness detectors to assess whether the simulated brains had transitioned to a conscious-like profile. In addition to the individual detector outputs, we computed a composite consciousness prediction by multiplying the cortical, thalamic, and pallidal detector scores, assigning equal weight to each.

As shown in Fig. 6, low-frequency stimulation (*<* 30 Hz) of thalamic projection nuclei modestly increased AI-predicted consciousness, consistent with clinical findings [18]. Moreover, as shown in Supplementary Fig. 8, the burstiness of the simulated thalamus at baseline negatively correlated with AI-predicted improvement in consciousness with 100-Hz DBS (*p* = 0.007), consistent with clinical data showing that patients with more tonic thalamic firing benefit most from intralaminar DBS at this frequency [19]. High-frequency (*>* 50 Hz) GPe stimulation also improved predictions in most simulated DOC cases, aligning with prior evidence [20]. However, the strongest and most consistent effects came from high-frequency stimulation of the STN, which significantly outperformed GPe stimulation in overall consciousness predictions (*p <* 0.001, Wilcoxon signed-rank), as well as in cortical (*p <* 0.001) and pallidal (*p <* 0.001) detectors (no advantage was seen in the thalamic detector, *p* = 0.5117). An example cortical LFP before and after STN stimulation is shown in Fig. 6E. STN effects were also robust across stimulation amplitudes (Supplementary Fig. 7).

**Fig. 6:**
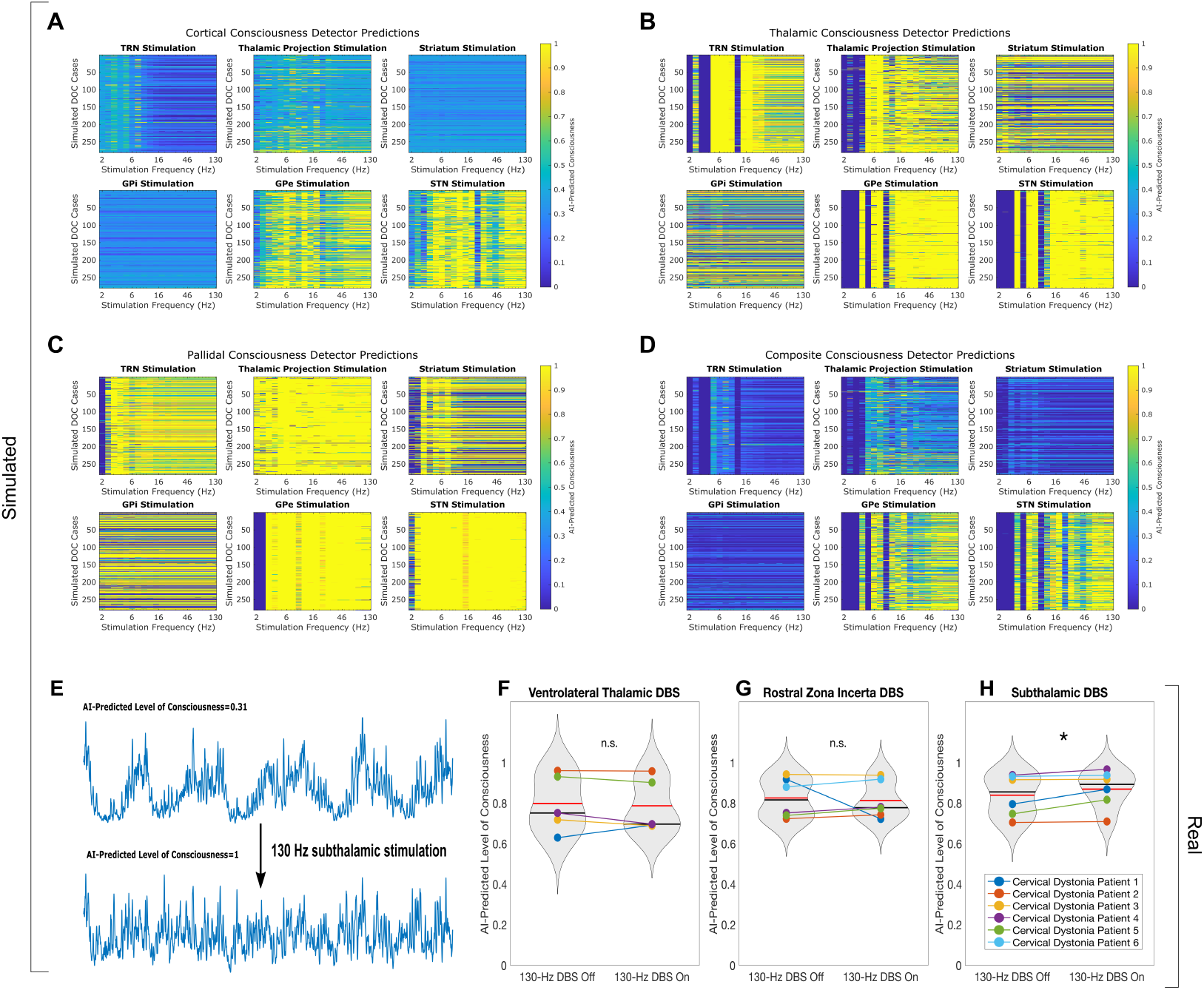
Effect of deep brain stimulation on AI-predicted levels of consciousness. **A–D**: AI-predicted consciousness levels across DBS parameters simulated in 280 model DOC cases. Deep brain stimulation was emulated using a square-wave input to subcortical targets in a biophysically grounded neural field model of the brain. Predictions from trained deep neural networks (DNNs) were used to assess the transition to conscious-like activity, evaluated across stimulation frequencies (2–130 Hz, logarithmic scale on x-axis) and DOC cases (y-axis). Each heatmap shows color-coded detector output for cortical (**A**), thalamic (**B**), and pallidal (**C**) DCNN-based consciousness detectors. Panel **D** shows the composite prediction, calculated by multiplying the cortical, thalamic, and pallidal outputs. High-frequency STN DBS (*>*50 Hz) elicited the most consistent increase in AI-predicted consciousness across detectors. **E**: Example cortical LFP trace from the simulated DOC model before and after high-frequency STN stimulation, with associated predictions from the cortical consciousness detector. **F–H**: Empirical validation of STN DBS predictions in real patients. We assessed 130-Hz DBS ON vs OFF in awake cervical dystonia patients implanted with electrodes targeting the ventrolateral thalamus (**F**), zona incerta (**G**), and SRN (**H**). For each patient, cortical EEG recordings were analyzed using the cortical DCNN consciousness detector. Violin plots show cross-patient distributions of AI-predicted consciousness for OFF and ON conditions, with connecting lines indicating per-subject mean trajectories. STN DBS significantly increased AI-predicted consciousness compared to baseline (*p* = 0.0156), consistent with simulation results. No significant effect was observed for zona incerta or ventrolateral thalamic DBS.

We next sought to test the predicted efficacy of high-frequency STN DBS in increasing level of consciousness in real human patients. With the exception of a single case report of sudden emergence from anesthesia with STN DBS [21], the relationship between STN DBS and consciousness has not been investigated, nor has STN

DBS been tested in patients with DOC. In the absence of such trials, we investigated whether high-frequency STN stimulation could increase cortical arousal—as measured by our DCNN cortical consciousness detector—when applied to awake humans. Specifically, we analyzed EEG recordings from six cervical dystonia patients who underwent alternating epochs of 130-Hz DBS ON and OFF. Each patient had been chronically implanted with DBS, with leads spanning the ventrolateral thalamus, the rostral zona incerta, and the STN.

When comparing AI-predicted level of consciousness across OFF and ON epochs, no effect was observed with DBS to the ventrolateral thalamus (Wilcoxon signed-rank test, one-tailed, *p* = 0.7812) or zona incerta (*p* = 0.2812). In contrast, STN DBS led to a significant increase in AI-predicted level of consciousness during ON periods compared to OFF (*p* = 0.0156, Cliff’s Δ = 0.222), consistent with the predictions of our AI model. Neither stimulation amplitude (LME, *t*(1185.0) = −0.12, two-tailed *p* = 0.9072) nor baseline trial order (LME, *t*(551.0) = −1.40, two-tailed *p* = −0.1617) had significant effects, indicating that the elevation in predicted consciousness was not attributable to gradual temporal drift or cumulative stimulation dose. These findings provide empirical support for the STN as a potential novel target for modulating consciousness-related brain dynamics.

## DISCUSSION

We introduce a new framework for uncovering the neural mechanisms of consciousness and its disorders, and identifying how DBS might restore awareness. Integrating large-scale electrophysiological data, deep learning, and neural field theory, we developed a generative AI model of conscious and unconscious brain states. This approach not only reproduces known neural signatures of DOC but also predicts previously unrecognized mechanisms of pathological unconsciousness.

Without being explicitly programmed to do so, the generative AI model recapitulates core features of the mesocircuit hypothesis, including reduced connectivity across corticothalamic, thalamocortical, corticostriatal, and striatopallidal pathways (Table 1) [5, 13]. The model also reveals compromised intra-thalamic coupling between projection and reticular nuclei, consistent with the thalamus’s sensitivity to ischemic and traumatic damage, even in the absence of direct insult [22–24]. Indeed, DOC patients show intra-thalamic white matter injury [25], and thalamic atrophy severity correlates with impaired consciousness [26]. Degeneration of reticular inhibition onto projection neurons—highlighted by our model—has also been observed in both cortical stroke [27] and TBI-induced coma [28].

Our model reproduces another key tenet of the mesocircuit hypothesis, namely reduced firing in cortical and thalamic projection neurons alongside increased pallidal activity, consistent with PET findings across DOC etiologies [29]. The model also recapitulates established neural correlates of consciousness, including complexity and entropy [11, 15], edge-of-chaos criticality [9, 11], alpha-band power [14], and thalamocortical low-to-high frequency information flow [9].

Beyond replication, our framework extends the mesocircuit hypothesis with mechanistic granularity. Rather than broadly implicating striatal dysfunction and the GPi, our model differentiates D1-from D2-expressing striatal neurons and teases apart GPe from GPi roles. Most strikingly, it predicts that unconsciousness partly stems from selective disconnection of D2 striatal neurons from the GPe—a prediction validated in DOC patients using diffusion MRI. Compared to MCS+/eMCS patients, those in a vegetative state showed reduced bilateral striatum–GPe streamline counts, and CRS-R scores correlated with left-side streamline counts. No difference was found in striatum–GPi streamlines, but vegetative patients showed elevated fractional anisotropy (FA) in left striatum–GPi tracts, consistent with our model’s prediction of increased D1–GPi coupling. This aligns with earlier work showing increased FA in striatopallidal tracts in DOC [30].

Our model also identified heightened reticular nucleus activity as a potential driver of unconsciousness, consistent with its role in generating cortical slow waves and suppressing arousal [31]. Though this has not been evaluated in DOC, increased TRN firing has been observed under anesthesia [32]. Simultaneously, the model predicted reduced reticular–thalamic coupling, in line with injury-related degeneration of inhibitory reticular inputs [27, 28]. We propose that this combination—hyperactive reticular firing alongside weakened connectivity—may impair thalamocortical gating, leading to fragmented modulation of cortical arousal and disrupted information flow (Table 1). Future work should test this mechanism empirically.

At the cortical microcircuit level, the model predicted increased excitatory-to-inhibitory and inhibitory-to-inhibitory coupling, along with reduced thalamic and pallidal input to cortical interneurons. Increased excitatory-to-inhibitory synaptic coupling has previously been observed in models of cortical injury [33]. While DTI studies confirm broad thalamocortical disconnection in DOC [34], they lack the resolution to isolate projections to inhibitory vs. excitatory neurons. Notably, our model predicts selective loss of subcortical input to inhibitory interneurons. This may explain why the presence of sleep spindles, which arise specifically from thalamic input to cortical interneurons [35], correlates with preserved awareness in DOC [36].

To test our model’s prediction of increased cortical inhibitory-to-inhibitory coupling in DOC, we reanalyzed single-nucleus RNA-seq data from two severe brain injury models: human coma after TBI and a rat model 48 hours post–middle cerebral artery occlusion. In both, we observed significant upregulation of VGF and SCG2 in parvalbumin-positive (PV^+^) cortical (or putative cortical) interneurons—the fast-spiking subclass modeled in our simulations. Notably, VGF is sufficient to drive PV^+^ *→*PV^+^ synaptogenesis in adult cortex [16], suggesting a conserved transcriptional response that enhances PV^+^ *→*PV^+^ cortical coupling after injury. At first glance, these findings may appear to contradict Nichols et al. [33], who reported reduced spontaneous IPSC frequency onto fast-spiking interneurons after cortical impact in juvenile mice. However, they found no change in IPSC amplitude or charge—arguing against broad synapse loss—and noted slowed IPSC kinetics, possibly reflecting receptor or dendritic changes. Moreover, because IPSCs onto PV interneurons originate from diverse sources, their results cannot be attributed specifically to cortical PV^+^ *→*PV^+^ connections. In contrast, our transcriptomic findings are cell-type and circuit-specific, identifying upregulation of genes shown to drive PV^+^ *→*PV^+^ synaptogenesis. These results raise the possibility that targeting the VGF/SCG2 axis could offer novel therapeutic avenues for reversing pathological cortical dynamics in DOC and severe brain injury.

Our generative AI model also revealed several novel, empirically untested correlates of unconsciousness. The strongest predictor was increased conduction velocity along axons connecting local cortical fast-spiking inhibitory interneurons—an unexplored mechanism in DOC or brain injury. However, increased PV^+^ interneuron synchrony during isoflurane anesthesia has been reported [37], which may reflect enhanced conduction along PV^+^ *→* PV^+^ axons. The model also predicted circuit-specific changes in EPSP duration, which remain to be tested empirically.

Critically, our framework offers a platform for in silico testing of DBS strategies in DOC. Across modeled cases, high-frequency (50–130 Hz) stimulation of the STN consistently increased predicted consciousness. This aligns with our observed increases in AI-predicted consciousness in cervical dystonia patients receiving high-frequency STN DBS. While direct evidence in DOC patients is lacking, multiple other findings support this target. In one intraoperative case, STN DBS triggered immediate return of consciousness and a surge in EEG entropy [21]. In rodents, STN stimulation after stroke enhances glucose metabolism in cortex and thalamus—regions often hypometabolic in DOC [38]—and drives sleep-to-wake transitions [39]. Moreover, STN-centered damage has been associated with abrupt induction of a vegetative state following radiofrequency exposure [40]. Together, these in silico, animal, and human data converge to identify the STN as a promising, yet unexplored, DBS target for restoring consciousness in DOC.

Several limitations remain. While the model was tuned with electrophysiology from cortex, thalamic projection nuclei, and pallidum—and validated against striatal recordings—it lacks direct empirical constraints for key regions like the reticular and subthalamic nuclei. Future work should incorporate recordings from these areas to enhance biological fidelity. The mean-field approach also abstracts away cellular diversity, notably failing to distinguish between matrix and core thalamic cells, which play distinct roles in consciousness [41, 42]. The model also does not resolve specific thalamic nuclei or cortical regions. Moreover, while our DCNN-based consciousness detectors offer partial interpretability—e.g., the cortical DCNN shows selectivity for sub-10 Hz and *β*-range features (Supplementary Fig. 1)—our approach embraces the DCNNs’ semi–black box nature as a strength. By learning directly from large-scale data, rather than relying on predefined features, these networks avoid investigator bias and generalize across species and modalities.

Further, while our snRNA-seq analysis revealed upregulation of synaptogenic genes in PV^+^ interneurons, this has not yet been confirmed histologically. Transcriptomic data provide indirect evidence and cannot quantify synapse number. Similarly, although our DTI analysis identified weakened striatum–GPe connectivity in DOC, it lacks cell-type specificity and cannot directly confirm synaptic loss. Validating these predictions will require histological studies of DOC brain tissue with region- and cell-type-specific resolution. Key therapeutic predictions of our model, such as the efficacy of subthalamic DBS, also await testing in animal models and clinical trials.

Finally, as AI tools gain influence in DOC science and care, their ethical implementation is critical. Transparency, explainability, and safeguards against overreliance on black-box models are essential—particularly in contexts of impaired communication and autonomy. Clinical oversight must remain central as these tools move toward real-world application.

While applied here to disorders of consciousness, the broader utility of this generative AI framework lies in its ability to model complex systems. Analogous approaches could be deployed to uncover mechanisms in other psychiatric or neurological disorders, or even non-neural systems characterized by hidden states and noisy observations. By uniting interpretable dynamical modeling with deep neural network discriminators, this platform may serve as a template for causal discovery in other scientific domains.

## Acknowledgments

We thank Diego Mateos for providing MEG data from a human epilepsy patient with absence seizures. We also thank Micah Johnson for help with deterministic tractography. This work was supported by NIH grant R01 GM135420, DOD CDMRP grant HT9425-24-1-1081, the Brain Injury Research Center at UCLA, the Fund for Scientific Research-Flanders (FWO), the Canada Excellence Research Chair in Neuroplasticity, the Belgian National Fund for Scientific Research, the fund Generet of King Baudouin Foundation, the European Foundation for Biomedical Research, and the Foundation for Research and Rehabilitation of Neurodegenerative Diseases.

## AUTHOR CONTRIBUTIONS

Conceptualization: DT, AB, NP, MM. Methodology: DT, ZSZ, JT, JG, AB, NP, MM. Investigation: DT, ZSZ, JT, JA, JG, HM, PV, CS. Data Curation: DT, ZSZ, JT, JA, JG, HM, KM, PV, CS. Supervision and Project Administration: AB, AH, SL, NP, MM. Writing and Editing: DT, JT, JG, HM, KM, PV, AB, NP, MM. Funding Acquisition: AH, NP, MM, SL, JA.

## Data Availability

Trained DCNN consciousness detectors, as well as configuration files to simulate our models of conscious and DOC brains, will be available on Figshare upon publication of this manuscript.

## METHODS

### Deep Neural Network Training

To generate a realistic computational simulation of conscious and comatose brain states, we first trained a deep convolutional neural network (DCNN) capable of accurately detecting consciousness from cortical electrophysiology recordings. The initial training data consisted of clinical EEG recordings from 22 ICU patients in acute coma from TBI (see below for patient demographics and EEG preprocessing). All neural data were z-score normalized prior to training. The neural network architecture comprised convolutional layers, batch normalization, ReLU activations, average pooling layers, and fully connected layers, concluding with a custom linear output layer designed to produce predictions bounded between 3 and 15, matching the scale of the Glasgow Coma Scale (GCS). This bounded output was then linearly remapped onto a standardized consciousness scale from 0 (fully unconscious) to 1 (fully conscious). Network hyperparameters—including learning rate, batch size, convolutional filter size, pooling stride, and the number of fully connected units—were optimized using Bayesian optimization. The final model was trained using the RMSprop optimizer with gradient clipping to maintain stable training dynamics. Validation was performed every 30 training iterations using 20% of the data, randomly set aside.

After initial training on acute coma EEG data, the cortical DCNN was further refined using cortical ECoG data from awake, fully conscious African green monkeys (*Chlorocebus aethiops*), as well as awake human essential tremor (ET) and Parkinson’s disease (PD) patients (see below for details on animal and patient data). To facilitate effective cross-species generalization, the learning rates for convolutional and fully connected layers were reduced by factors of 0.05 and 0.25, respectively. Subsequently, the network was refined again, simultaneously incorporating EEG and ECoG datasets, ensuring robust generalization across species and recording modalities.

We then incorporated EEG recordings from 14 patients with chronic disorders of consciousness (DOC), including vegetative and minimally conscious states (see patient demographics below). These patients’ consciousness levels were clinically evaluated using the Coma Recovery Scale-Revised (CRS-R). To integrate CRS-R scores with the unified 0-to-1 consciousness scale, we first performed a piecewise linear mapping from CRS-R scores onto the GCS scale, followed by the linear remapping described above. Specifically: CRS-R scores from 0–5 were linearly mapped to GCS scores of 3–5, corresponding to consciousness levels from 0.00 to 0.167; CRS-R scores from 6–10 were mapped to GCS scores of 6–8, corresponding to consciousness levels from 0.25 to 0.417; CRS-R scores from 11–15 were mapped to GCS scores of 9–11, corresponding to consciousness levels from 0.50 to 0.667; and CRS-R scores from 16–23 were mapped to GCS scores of 12–15, corresponding to consciousness levels from 0.75 to 1.00. This continuous, piecewise linear mapping ensured smooth integration of chronic DOC data within our standardized consciousness framework, capturing subtle variations in patient consciousness accurately. Finally, the network underwent additional fine-tuning (using previously reduced learning rates) on resting-state EEG data from healthy, awake human controls (Dataset 1, see below), reinforcing its ability to reliably detect fully conscious states.

We also trained a second DCNN to discriminate real cortical electrophysiological signals from synthetic signals. Synthetic datasets included 500 amplitude-modulated signals, 500 frequency-modulated signals, 500 sawtooth waves, 500 square waves, 500 parametric sine waves, 500 Lorenz-system simulations, and 500 instances each of pink, blue, and red noise, each created with randomly varied parameters. This synthetic-versus-real network shared a similar architecture with the consciousness-detection network, concluding with a bounded output layer for binary classification. Real electrophysiology data for training included EEG from human coma patients, ECoG from monkeys, human PD and ET patients, epilepsy patients, and MEG from a human epilepsy patient. Data were z-score normalized, balanced between synthetic and real data, and the network was trained using the RMSprop optimizer with Bayesian hyperparameter optimization, gradient clipping, and periodic validation every 30 iterations.

Additionally, a third DCNN was trained to differentiate generalized seizure data from non-seizure data. Non-seizure recordings included normal waking data from humans and rats with epilepsy, ECoG from human ET and PD patients, monkey recordings during wakefulness and anesthesia, and EEG from acute coma patients across the full GCS range. Generalized seizure data included previously published human ECoG and MEG recordings, supplemented by generalized seizure and normal waking data from seven Genetic Absence Epilepsy Rats from Strasbourg (GAERS) - see below for details on data acquisition and preprocessing. The network architecture mirrored previous models, optimized via Bayesian hyperparameter tuning, and utilized RMSprop with gradient clipping and periodic validation.

Two further DCNNs were trained to classify conscious versus anesthetized states from subcortical local field potentials (LFPs), one for thalamic data (from human ET patients and Long-Evans rats) and one for globus pallidus externa (GPe) data (from human PD patients and two African green monkeys). These networks also followed similar architectures and optimization protocols, employing Bayesian hyperparameter optimization, RMSprop training, gradient clipping, and periodic validation.

Outputs from these trained DCNNs provided the loss function foundation for a genetic optimization algorithm, which tuned parameters of a biophysically realistic mean-field model of the basal ganglia-thalamo-cortical system (described below). After identifying model parameters that initially simulated conscious states—confirmed by predictions close to 1 (fully conscious), non-synthetic, and non-seizing cortical dynamics, alongside conscious thalamic and pallidal classifications—we trained three additional discriminator DCNNs. These discriminators distinguished simulated cortical, thalamic, and pallidal activity from corresponding real recordings during conscious states. For training, we generated 1,000 simulated 10-second segments per region, divided into training (800) and validation (200) subsets. Equivalent numbers of real data segments were selected from previous conscious-state recordings for balanced training and validation. Network architectures, hyperparameter optimization, training strategies, and validation procedures were consistent with previous models. These discriminators were subsequently employed to further enhance the realism of our computational brain simulations, as detailed below.

### Phase-Amplitude Coupling Analysis

Phase-amplitude coupling (PAC) patterns corresponding to waking, conscious states were identified within the cortex, thalamus, and GPe, as well as between cortex-thalamus and cortex-GPe in both directions. For the thalamic and thalamo-cortical analyses, we used all available data from human ET patients and Long-Evans rats. For the GPe and pallido-cortical analyses, data from human PD patients and African green monkeys were used. The intra-cortical analysis utilized data from human ET and PD patients, Long-Evans rats, and African green monkeys. PAC was computed by analyzing the modulogram from each 10-second segment or sample of data. To do so, we first filtered the source and target LFP signals to extract phase and amplitude across a range of frequencies (in the case of intra-regional PAC, the source and target signals were the same). Phase frequencies spanned 1 to 15 Hz, while amplitude frequencies ranged from 25 to 105 Hz. The phase of the source signal and the amplitude envelope of the target signal were computed using Hilbert transforms, and the modulation index (MI) was calculated within predefined phase bins. For each animal or patient, we computed a mean modulogram across all data segments, resulting in an average PAC profile for each individual. These individual profiles were then averaged across animals or patients to generate overall modulograms for intra-cortical, intra-thalamic, intra-pallidal, cortico-thalamic, thalamo-cortical, cortico-pallidal, and pallido-cortical PAC. These empirical PAC patterns were then incorporated into the loss function of our genetic optimization algorithm, to create a simulation of the conscious brain with realistic PAC.

### Mean-field model and genetic optimization

In this study, we employed a mean-field model similar to the one described in previous work to simulate LFPs and the average firing rates of various neural populations [1–3], based on the original thalamo-cortical-basal ganglia neural field model described by van Albada and Robinson [2]. The model incorporates several biophysically grounded processes including a nonlinear sigmoidal firing response, dendritic and somatic filtering, finite axonal conduction delays, and spatial propagation of activity via a damped wave mechanism.

The firing rate, denoted as *Q*_*a*_(*r, t*), represents the population-average proportion of neurons within population *a* whose membrane potential exceeds their reversal potential. This is modeled as a sigmoidal function of the mean membrane potential *V*_*a*_ relative to the mean-field threshold *θ*_*a*_ (mV) at which the population’s firing rate reaches half its maximum:

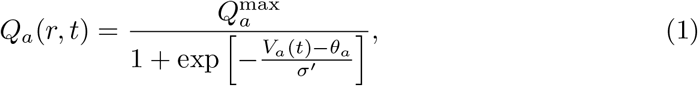

where 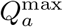 is the maximum possible firing rate and *σ*^*′*^ is the standard deviation of the subthreshold membrane potentials around *θ*_*a*_.

The change in the mean membrane potential *V*_*a*_(*t*) over time is determined by the combined synaptic input from other neural populations. Each input undergoes dendritic filtering that is well approximated by a convolution with a response kernel that embodies a fast rise and subsequent exponential decay. By approximating this convolution with a differential operator (after appropriate normalization by *η*(*α, β*)), we express the membrane potential dynamics as

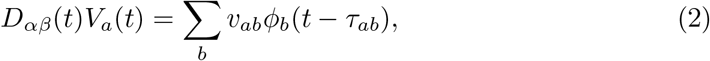

where *v*_*ab*_ is the axonal-synaptic coupling strength from population *b* to *a, ϕ*_*b*_(*t* − *τ*_*ab*_) is the firing rate arriving from population *b* with a delay *τ*_*ab*_ (which captures finite axonal conduction and synaptic transmission delays), and *D*_*αβ*_(*t*) is the differential operator given by

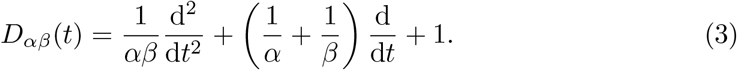

Here, *α* and *β* represent the decay and rise rates of the dendritic/somatic response, respectively. Note that this differential operator is derived as an approximation to the convolution of incoming signals with the dendritic response kernel *L*_*ab*_(*t*); the factor *η*(*α, β*) in later equations ensures proper normalization.

The synaptic response itself was modeled as

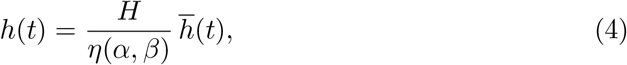

where 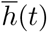 is the original synaptic impulse response and *H* is a synaptic scaling factor. In contrast to our previous work [1] where *H* was fixed at 31.5 s^−1^, here *H* is allowed to vary independently for each brain region.

The outgoing mean electric field, *ϕ*_*ab*_(*r, t*), which represents the axonal propagation of activity from population *b* to *a*, is modeled either using a damped wave equation or, for certain connections, via a direct mapping. In the original van Albada and Robinson model [2], connections expected to exhibit wave-like propagation are described by

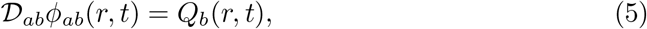

with the differential operator

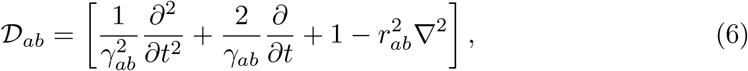

where *r*_*ab*_ is the spatial range of axonal projections, *γ*_*ab*_ is the temporal damping coefficient, and *∇*^2^ is the Laplacian operator. This formulation arises as a two-dimensional generalization of the telegrapher’s equation, and in the limit of minimal spatial variations (or for local neural mass approximations), the Laplacian term becomes negligible so that the operator reduces to that of a damped harmonic oscillator. In contrast, connections that were originally modeled as simple maps in van Albada’s model [2] are implemented using a direct mapping, i.e.,

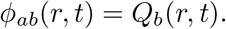

All outgoing connections from the GPe introduced in this and our prior modification of the model [1] are direct mappings.

Time delays *τ*_*ab*_ are explicitly included in the formulation to account for both finite axonal conduction speeds and synaptic transmission delays, and they play a crucial role in shaping the dynamic, oscillatory, and wave-like behavior of the model. Finally, external inputs to the system were modeled as white noise applied to the thalamic projection nucleus, representing stochastic background activity or sensory noise that drives the system. The circuit connectivity was based on the original van Albada and Robinson model [2], with additional projections from the GPe to cortical and thalamic regions incorporated as in our recent modifications [1]. In the model, intralaminar and relay nuclei are collectively treated as a “projection” nucleus. Likwewise, the brainstem and sensory organs are collectively treated as a single region, the activity of which is modeled as a random univariate Gaussian signal. In our simulations, each population was assigned a length of 10 cm but was modeled using a single node, thereby neglecting any within-population spatial variation. We used a fixed time-step of Δ*t* = 0.0001 for numerical integration. For the wave propagators, integration was performed using an explicit finite-difference approach, which specifies a nine-point spatial stencil to reduce high-frequency instabilities when the system is driven by a random Gaussian process.

Model parameters were optimized using a genetic algorithm, similar to our previous work [1], to identify physiologically realistic configurations for the conscious brain. Genetic algorithms are well-suited for optimizing multiple objectives and navigating complex, non-convex parameter spaces, and allow for interpretable solutions without requiring gradient information.

The fitness function integrated multiple empirical constraints and DCNN outputs. Specifically, we required that (1) all regional firing rates fell within physiologically observed ranges; (2) fluctuations in cortical firing rates were positively correlated with high-gamma (60–200 Hz) LFP spectral power [4]; (3) cross-frequency information transfer between the cortex and thalamus reproduced empirically observed spectral asymmetries [1]; (4) phase-amplitude coupling (PAC) patterns within and between the cortex, thalamus, and GPe matched those observed in real data; and (5) DCNN classifiers identified the simulated cortex, thalamus, and GPe as consistent with consciousness, detected no seizure-like activity in the cortex, and failed to distinguish the simulated cortical data from real human recordings.

In addition to these constraints, we required that the cortical LFP dynamics exhibit weak chaos, as measured by logarithmic divergence of nearby trajectories, consistent with prior evidence that conscious cortical electrodynamics lie near edge-of-chaos criticality [1, 5]. No such constraint was imposed on subcortical LFPs.

The composite fitness function was defined as:

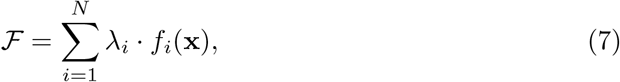

where **x** denotes the model parameter vector, *f*_*i*_ is the *i*-th objective (e.g., firing rate error, information transfer asymmetry, classifier output, chaos score), *λ*_*i*_ is the corresponding weight, and *N* is the total number of components. The weights *λ*_*i*_ were manually adjusted between optimization batches to emphasize underperforming features—for instance, increasing the penalty for low cross-frequency transfer or poor classifier agreement.

Each batch consisted of 50 parallel genetic algorithm runs. At the end of each batch, we manually reviewed the resulting simulations and selected the best-performing parameter vectors as seeds for the next round. Over 118 such optimization batches, we gradually refined the solution space until a single parameter set was found that satisfied all empirical and DCNN-based constraints simultaneously.

At that point, we trained three additional discriminator networks to classify simulated versus real recordings from the cortex, thalamus, and GPe. These discriminators were then incorporated into the fitness function as adversarial components, penalizing model parameters that produced outputs distinguishable from real conscious brain activity in any of the three regions. We then resumed optimization—again running 50 simulations in parallel per batch, manually adjusting weights between batches, and selecting only the top-performing simulations as seeds. After 76 additional optimization batches, a single parameter vector was found that satisfied all criteria, including the adversarial discriminator networks. This parameter set was used as the final model of the conscious brain state. All parameter values are listed in Tables S2–S4.

### TBI acute coma EEG recordings (training and validation data)

EEG data from 22 patients in acute coma due to TBI (mean age 31.5 *±* 11.3 years, 18 males, 4 females) were collected at the UCLA Ronald Reagan Medical Center and used to train the cortical consciousness detector. These data have been described in detail in a previous paper from our group [6]. All data were sampled at 250 Hz and bandstop filtered between .5 and 45 Hz. Patients were included based on GCS scores *<*9 or GCS 9-14 with CT evidence of intracranial bleeding. Patients with non-significant head injuries, prior neurological disease, or brain death were excluded. Patients’ conscious state was assessed multiple times daily using the GCS. EEG segments were spaced at least 12 hours apart. Data were divided into 10-second trials, cleaned for line noise with a band-stop filter, detrended, demeaned, and re-referenced to the common average. Independent component analysis (ICA) was then performed to remove ocular and cardiac artifacts, and samples with significant motion artifacts were excluded after visual inspection.

### Chronic DOC EEG recordings (training and validation data)

EEG data from fourteen patients with chronic disorders of consciousness (0.97-6.8 years since injury) resulting from mixed etiologies (traumatic or non-traumatic injuries) (mean age 38.5 *±* 16.3 years, seven females, seven males) were used to train the cortical consciousness detector. Data were recorded at Casa Colina Hospital and Centers for Healtchare as part of an ongoing study on the effect of low-intensity focused ultrasound on awareness in chronic DOC. The study was approved by Institutional Review Boards of Casa Colina Hospital and Centers for Healtchare and UCLA. Continuous EEG data were recorded at 1,000 Hz at baseline (before administration of low-intensity focused ultrasound) and subsequently resampled to 250 Hz and bandstop filtered between 0.5 and 45 Hz to match the acute TBI EEG data. EEG recordings were divided into 10-second samples, band-stop filtered for line noise and harmonics, detrended, demeaned, and re-referenced to the common average. ICA was then performed to remove cardiac and ocular artifacts, and samples with significant motion artifacts were excluded after visual inspection.

### *Chlorocebus aethiops* anesthesia ECoG recordings (training, validation, and independent test data)

Simultaneous intracranial electrophysiology recordings from the frontal cortex and GPe of African green monkeys (*Chlorocebus aethiops*) were reused from a previously published study [7] to train the pallidal consciousness detector and evaluate cortical and pallidal phase-amplitude coupling patterns. The study, approved by The Hebrew University (MD-18-15449-5), involved two surgeries under deep anesthesia. In the first, a recording chamber and EEG electrodes were implanted to target the cortex and GPe. In the second, a venous catheter was inserted. MRI confirmed correct placement of the recording chamber, and recording sessions followed a protocol alternating between baseline, sedation, and washout, with vital signs monitored. During recordings, microelectrodes captured raw signals, spikes, and LFPs at a sampling frequency of 1375 Hz. To match the cortical EEG data, cortical ECoG signals were resampled to 250 Hz and bandstop filtered between 0.5 and 45 Hz; subcortical LFP signals were resampled to 500 Hz with no bandstop filtering. Here, only recordings during waking and propofol anesthesia states were used. Recordings were divided into 10-second samples, detrended, demeaned, and re-referenced to the common average. Line noise was removed with a band-stop filter, and samples with significant motion artifacts were excluded. Data were provided by J.G.

### Parkinson’s disease patient ECoG and LFP anesthesia recordings (training and validation data)

Data from a prior study from our group [8], which included pallidal LFP and frontoparietal ECoG data from 12 PD patients who underwent GPi DBS lead implantation, were reused here to train our pallidal consciousness detector and evaluate pallidal-cortical phase-amplitude coupling. The average age of the patient cohort was 65 *±* 7 years, and included three female patients. All patients provided informed consent, and PD medications were stopped 12 hours prior to surgery. Preoperative MRI and postoperative CT scans were used to target the motor GPi for lead placement, confirmed via intraoperative stimulation testing. An 8-contact ECoG strip was temporarily inserted through the frontal burr hole for cortical and depth electrode LFP recordings (from both GPe and GPi) after DBS implantation and propofol administration.

LFPs were recorded from the DBS lead’s 4-ring contacts (ventral to dorsal), with data acquired using BCI2000 and sampled at 2400 Hz. Recordings began with patients awake and resting before propofol was administered to induce loss of responsiveness. Data were split into the first minute waking pre-anesthetic bolus, and the last minute of recordings, by which point all subjects were verbally unresponsive. Localization of ECoG strip contacts in M1, S1, and DBS contacts was based on either postoperative imaging or the median nerve SSEP technique. Recordings were divided into 10-second segments, detrended, demeaned, band-stop filtered for line noise, and re-referenced to the common average, and visually inspected for segments with significant motion artifacts, which were excluded. ECoG data used for the cortical consciousness detector training were resampled to 250 Hz and bandstop filtered between 0.5 and 45 Hz (to match the other datasets), while the pallidal LFP recordings were resampled to 500 Hz without bandstop filtering.

### Healthy, awake human EEG recordings, dataset 1 (training and validation data)

EEG data from ten healthy, right-handed volunteers (six females, four males; mean age 30.25 *±* 6.95 years) were obtained from an open-source dataset previously described by Torkamani-Azar and colleagues [9]. Participants had normal or corrected-to-normal vision and were not taking any drowsiness-inducing medications. Following consent per Sabanci University’s Research Ethics Council, EEG data were recorded at a 256 sampling frequency in a dimly lit, Faraday-caged room. Monopolar EEG signals were acquired using 64 Ag/AgCl active electrodes placed according to the 10-10 system. Data from a 2.5-minute resting-state session with eyes open were used as part of the training for our cortical consciousness detector. To match the coma/DOC EEG used to train the cortical consciousness detector, recordings were resampled to 250 Hz and bandstop filtered between 0.5 and 45 Hz. Data were then divided into non-overlapping 10-second samples, filtered for line noise and harmonics, detrended, demeaned, and re-referenced to the common average. ICA was performed to remove cardiac and ocular artifacts, and segments with motion artifacts were removed following visual inspection.

### Human essential tremor patient ECoG and LFP anesthesia recordings (training and validation data)

Data from another previous study by our group [10] were re-analyzed to train the thalamic consciousness detector and assess thalamic-cortical phase-amplitude coupling. Recordings were collected from 10 ET patients (6 female, 4 male, ages 60–79) undergoing unilateral (n=6) or bilateral (n=4) DBS implantation in the ventral intermediate (VIM) nucleus of the thalamus. All participants gave written informed consent, and the study was approved by the UCLA institutional review board. LFPs were recorded from the VIM thalamus, and ECoG signals from the ipsilateral frontoparietal cortex during rest and following intravenous propofol administration. Signals were acquired at a 2400 Hz sampling rate and filtered between 0.1 and 1000 Hz.

Patients remained awake with eyes open for the first minute of recording, which was used as the “awake” state. Propofol was then administered until patients reached a MOAA/S score of 0 (no response) or 1 (response to noxious stimuli). LFP and ECoG recordings continued for 5 minutes after anesthesia, with the final minute used as the “anesthetized” state. Data were segmented into 10-second segments, demeaned, detrended, and line noise was removed using a band-stop filer. Segments with multi-channel artifacts were excluded after visual inspection. For the cortical consciousness detector training, ECoG data were resampled to 250 Hz and bandstop filtered between 0.5 and 45 Hz (this filtering was done to ensure compatibility with the other datasets). The thalamic LFP recordings, however, were only resampled to 500 Hz.

### Healthy, awake human EEG recordings, dataset 2 (independent test data)

An independent test dataset of EEG recordings from 20 healthy adult participants (16 males, 4 females; age range: 21–31 years) was obtained from the Auditory Evoked Potential EEG-Biometric dataset v1.0.0 [**?**], publicly available via PhysioNet [11]. All recordings were conducted at the Faculty of Engineering of Marche Polytechnic University in Ancona, Italy, and followed the ethical guidelines of the World Medical Association Declaration of Helsinki. All participants gave informed consent prior to participation.

In the present study, we used only the eyes-open resting-state EEG recordings. Each subject completed three recording sessions in a single day, with three minutes of resting-state, eyes-open EEG collected during each session. Data were acquired using an OpenBCI Ganglion board at a sampling rate of 200 Hz, with gold cup electrodes placed at four scalp positions (T7, F8, Cz, P4) in accordance with the 10/10 international EEG system. Reference and ground electrodes were placed at the left and right ears, respectively.

For preprocessing, data were divided into non-overlapping 10-second segments or samples, resampled to 250 Hz (using piecewise cubic Hermite interpolation to generate intermediate samples), cleaned for line noise with a band-stop filter, detrended, demeaned, and re-referenced to the common average. ICA was then performed to remove ocular and cardiac artifacts, and samples with significant motion artifacts were excluded after visual inspection.

### Cardiac arrest acute coma EEG recordings (independent test data)

EEG data from comatose ICU patients following cardiac arrest were obtained from the I-CARE (International Cardiac Arrest REsearch consortium) Database v2.1 [12, 13], publicly available on PhysioNet [11]. This multi-center dataset includes clinical and continuous EEG recordings, sampled at multiple frequencies, from 607 adult patients across seven academic hospitals in the United States and Europe. All patients had return of spontaneous circulation (ROSC) after in- or out-of-hospital cardiac arrest but remained comatose, defined as a Glasgow Coma Score (GCS) *≤* 8 and inability to follow verbal commands. Continuous EEG was typically initiated within hours of ROSC and continued for hours to days.

For this study, only each patient’s first comatose EEG dataset was used. Recordings were provided in WFDB-compatible format and varied in duration, channel configuration, and recording quality across hospitals and patients. To ensure consistency, we excluded EEGs with fewer than 60 seconds of usable data following preprocessing.

Preprocessing was performed using the FieldTrip toolbox in MATLAB. EEG recordings were segmented into non-overlapping 10-second samples, downsampled to 250 Hz, and bandpass filtered between 0.5 and 45 Hz. All data were further bandstop filtered to remove utility-line harmonics and detrended and demeaned. To mitigate artifact contamination, we excluded the first 20 seconds of every recording and used a multi-stage rejection pipeline. First, we implemented the “Twin Peaks” algorithm for spike-and-wave seizure detection [14], removing all samples containing spike-wave discharges. Second, we computed the standard deviation of the EEG signal for each channel and data segment, rejecting segments and channels with outlier standard deviations (threshold: *>* 2.5 SD from the mean). Channels with persistently high standard deviation across segments were removed entirely. Third, independent component analysis (ICA) was applied to the remaining data, and components were visually inspected to identify residual epileptiform activity, ocular artifacts, cardiac artifacts, or other artifactual sources. Recordings in which most ICA components appeared pathological or artifactual were excluded in full.

Only recordings with at least 60 seconds of post-cleaning data were retained. Across the final dataset, 455 patients met all inclusion criteria and were included as part of the independent test evaluation for the cortical consciousness detector.

### Long-Evans rat anesthesia ECoG and LFP recordings (training, validation, and independent test data)

Previously published data from Reed and Plourde [15] were reanalyzed to train a thalamic consciousness detector. Bipolar electrodes were placed in the ventral posterior nucleus of the thalamus and ipsilateral sensory cortex, with a reference electrode in the parietal bone. Propofol was administered intravenously, achieving plasma concentrations of 3, 6, 9, and 12 *µ*g/ml. LFPs were recorded at a sampling frequency of 3030 Hz after 15 minutes of drug equilibration, with unconsciousness (loss of righting reflex) occurring at 9 *µ*g/ml. LFPs from awake and 12 *µ*g/ml conditions were used in this study. Signal were divided into 10-second segments, demeaned, detrended, band-stop filtered to remove line noise, and visually inspected for motion artifacts. Cortical recordings, used as independent test data for the cortical consciousness detector, were downsampled to 250 Hz and bandpass filtered between 0.5 and 45 Hz. Thalamic LFPs, which were used as training and validation data for the thalamic consciousness detector, were resampled to 500 Hz without bandpass filtering.

### Human and rat ECoG, EEG, and MEG generalized seizure data

Data from Miyamoto and colleagues [16] were reanalyzed to train the generalized seizure detector. Seven GAERS rats with spontaneous 6-8 Hz spike-and-wave seizures were implanted with stainless steel ECoG electrodes in the somatosensory cortex and a reference electrode on the cerebellum. LFPs were simultaneously collected from their dorsal striatum. Only seizures lasting at least 10 seconds were used. The data were split into 10-second segments, demeaned, detrended, and filtered at line noise and harmonics, with segments containing multi-channel artifacts removed. Seizures were manually labeled based on LFP traces.

In addition, data from two human epilepsy patients with secondarily generalized seizures, downloaded from the European Epilepsy Database [17] and described in detail in our prior study [5], were also used to train the generalized seizure detector. Subject 1, a 42-year-old male with cortical dysplasia, had six electrode strips (26 electrodes) over the right temporal cortex. Subject 2, a 14-year-old female with cryptogenic epilepsy, had one grid and six strips (96 electrodes) over the left temporal cortex. ECoG signals were recorded at 1024 Hz, resampled to 500 Hz, detrended, and re-referenced. Seizures lasting at least 10 seconds were selected, and artifact-laden electrodes were removed. Additionally, MEG data from an 18-year-old female with absence seizures [18] were used. The data, recorded at 625 Hz, were band-stop filtered for line noise, resampled to the common average, and cleaned using ICA to remove ocular and cardiac artifacts.

### Bat LFP recordings

We analyzed a previously published dataset of LFP recordings from the dorsal striatum of Seba’s short-tailed bats (*Carollia perspicillata*) during spontaneous communication calls [19]. The dataset was downloaded from the authors’ public repository [20] and consisted of 788 trials of striatal LFPs, each spanning 1001 samples (−500 to +500 ms relative to vocal onset) across three tetrode channels, originally sampled at 20 kHz and downsampled to 1 kHz. For the current analysis, we further downsampled the LFPs to 500 Hz. To facilitate comparisons with simulated data, we grouped the trials into non-overlapping sets of 10 and concatenated them sequentially along the temporal axis. This preprocessing enabled spectral power and phase–amplitude coupling analyses over extended temporal windows that matched the structure of the simulated signals. All surgical and recording procedures were approved by the Regierungspräsidium Darmstadt, Germany (permit number: FU1126). LFPs were recorded using a chronically implanted tetrode mounted on a microdrive, targeting the dorsal striatum, with signals digitized at 20 kHz using a portable multichannel system (Multi Channel Systems MCS GmbH).

### Single-nucleus RNA-seq Analysis in Human and Rat Cortex

To test model-predicted changes in cortical inhibitory-to-inhibitory synaptic coupling in disorders of consciousness (DOC), we reanalyzed publicly available single-nucleus RNA sequencing (snRNA-seq) datasets from both human patients in coma and rats subjected to severe ischemic stroke. In both species, we focused exclusively on parvalbumin-positive (PV^+^) inhibitory interneurons—the same fast-spiking subclass modeled in our AI-based simulations (mean firing rate *∼*18 Hz; Table S5)—and quantified expression of genes linked to PV *→*PV synapse formation and inhibitory synapse structure.

Human cortical snRNA-seq data were obtained from Garza et al. [21] (GEO accession GSE209552), including five healthy control donors and six patients in acute coma following traumatic brain injury (TBI). Nuclei were isolated from resected cortical tissue and processed using 10x Genomics Chromium Single Cell 3’ protocols. Reads were aligned to the GRCh38 genome using Cell Ranger v5 with intronic reads retained (`--include-introns`). Cell filtering and clustering were performed in Seurat v3.1.1 for R v3.4. From the processed gene-by-cell matrices, we extracted PV^+^ cells by gating on non-zero *PVALB* expression. Only donor samples containing PV^+^ cells were retained, yielding 5 control and 6 TBI samples for analysis. Gene expression was binarized at the single-cell level (expressed = any UMI), and generalized linear mixed-effects models (GLMMs) were fit with a binomial distribution and donor as a random intercept. For each gene, we tested whether PV^+^ cells from coma patients were significantly more likely to express the transcript compared to controls using a one-tailed test. Visualization of *VGF* and *SCG2* expression across the PV^+^ transcriptional manifold was performed using t-distributed stochastic neighbor embedding (t-SNE) on log_2_(CPM+1) data from principal components (perplexity 30), with expression values mapped to dot color, size, and transparency as described in the figure caption (Fig. 4A, D).

Rat snRNA-seq data were obtained from a recent study of ischemic stroke [22] (GEO accession GSE250245), which included three brain samples: right hemisphere of a sham-operated control, and ipsilateral (right) and contralateral (left) hemispheres from a rat 48 hours after severe permanent middle cerebral artery occlusion. All nuclei were isolated and processed using the 10x Genomics Chromium 5’ v2 workflow and sequenced on a NovaSeq 6000. Cell Ranger v7.0.0 was used for demultiplexing and alignment to the mRatBN7.2 genome. To isolate PV^+^ cortical interneurons, we retained cells expressing the canonical marker *Parva*, co-expressing *SATB1*, and lacking expression of subcortical markers *PTHLH* and *C1QL1*. These gating criteria excluded non-cortical PV^+^ subtypes and ensured anatomical specificity. For each gene, a GLMM with binomial distribution and donor as random intercept was used to assess the binary expression outcome across groups. Contrasts of interest included ipsilateral vs. sham and ipsilateral vs. contralateral cortex. Visualizations of *VGF* and *SCG2* expression were generated using t-SNE projection on the combined dataset of cortical PV^+^ cells, colored and scaled according to gene expression as in the human analysis (Fig. 4D).

Across species, we analyzed seven genes: the neuropeptide-encoding genes *VGF* and *SCG2*, which are both necessary and sufficient to drive PV^+^ *→*PV^+^ synaptogenesis in the adult cortex [23], and five canonical markers of inhibitory synapse structure: *ARHGEF9, GABRA1, GABRB2, GABRB3*, and *GPHN*. Expression results were quantified at both the single-cell level via GLMMs and the donor/sample level as the proportion of PV^+^ cells expressing each gene. The latter were plotted with standard error bars and per-donor scatter points for visual clarity (Fig. 4B–C, E–F; Fig. S5).

### DOC patient diffusion tensor imaging data

We analyzed structural and diffusion MRI data from 51 DOC patients recruited at the University of Liège. Diagnoses were made using the Coma Recovery Scale–Revised (CRS-R) [24]. The final dataset included 21 patients in a vegetative state (VS), 16 in a minimally conscious state with some preserved capacity for communication or command following (MCS+), and 13 in emergence from MCS (EMCS). Four patients in MCS-were excluded due to insufficient sample size. The cohort comprised 30 males and 20 females (mean age = 44.42 *±* 15.77 years). Etiologies were anoxic (n=12), traumatic (21), hemorrhagic (3), metabolic (1), infectious (1), and mixed/other (12). At the time of MRI acquisition, patients ranged from 8 days to 8.33 years post brain injury. Written informed consent was obtained from each patient’s legal guardian. The study was approved by the Ethics Committee of the University Hospital of Liège.

MRI data were acquired using a 3T Siemens Tim Trio scanner. T1-weighted images were collected with the following parameters: 120 slices, TR = 2300 ms, TE = 2.47 ms, voxel size = 1 *×* 1 *×* 1.2 mm, flip angle = 9°. Diffusion-weighted images (DWI) were acquired with 64 non-collinear directions at a b-value of 1000 s/mm^2^, repeated twice (45 axial slices, TR = 5700 ms, TE = 87 ms, voxel size = 1.8 *×* 1.8 *×* 3.3 mm, flip angle = 90°), along with one b=0 image. DWI data were denoised, corrected for motion and eddy currents, and checked for artifacts using DTIPrep [25]. Brain extraction was performed with BET [26] for diffusion images and optiBET [27] for T1-weighted images.

Three basal ganglia ROIs (striatum, external globus pallidus [GPe], and internal globus pallidus [GPi]) were taken from the Keuken and Forstmann probabilistic atlas [28], transformed from MNI to subject T1 space using ANTs [29], and then into diffusion space using FLIRT [30]. Patients were excluded if they exhibited a level of brain deterioration that obscured the delineation of the ROIs (e.g., registration failure during ROI transformation to subject space). CSF masks were generated and intersected with all ROIs to eliminate spurious voxels. All masks were visually inspected and manually refined as necessary.

Probabilistic tractography was performed in native diffusion space using FSL’s `probtrackx2`, modeling up to three fiber orientations per voxel. For each hemisphere, 5000 streamlines were launched from every voxel in the striatal seed mask. Target masks were defined as the external and internal globus pallidus (GPe and GPi, respectively), with the GPi serving as a stop mask during tractography to the GPe, and the GPe serving as a stop mask during tractography to the GPi. This approach ensured that streamlines terminating in the target did not pass through the alternate pallidal segment, improving anatomical specificity.

To quantify structural connectivity, we computed the total number of streamlines that reached each target mask. To account for anatomical variability across patients, we also computed a geometric normalization of the streamline counts by dividing the raw count by the square root of the product of the seed and target mask sizes (i.e., 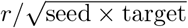). This geometric mean normalization mitigates bias introduced by inter-individual differences in ROI volume while preserving biologically meaningful variation in connectivity strength. To assess microstructural properties along the tracts, we extracted the mean fractional anisotropy (FA) for tracts to each pallidal ROI. FA, a scalar derived from the eigenvalues of the diffusion tensor, reflects the directional coherence of water diffusion and serves as a widely used index of white matter integrity, with higher values typically indicating greater anisotropy and fiber organization [31]. All metrics were computed separately for the left and right hemispheres.

To generate the example deterministic tractography visualizations in Fig. **??**, we used DSI Studio. As in our probabilistic tractography analysis, the striatum served as the seed and the GPe as the target mask. Streamlines were permitted to pass through the target. Tracking was conducted in each subject’s native diffusion space without voxel resampling. Streamlines were generated using up to one million randomly placed seeds per subject, with no fractional anisotropy, curvature, or step-size thresholds imposed. Streamline length was constrained to a minimum of 5 mm and a maximum of 20 mm. Resulting tracts were overlaid on the subject’s own T1-weighted anatomical image for visualization, with striatal, GPe, and tractography masks rendered in pink, green, and blue, respectively.

### Cervical dystonia patient DBS data

We reanalyzed data originally collected and reported in Wiest et al. (2022) [32], which included EEG recordings from six patients (four female, two male; mean age 57.17 *±* 7.73 years) with isolated idiopathic cervical dystonia who underwent dual-targeted DBS of the STN and ventrolateral thalamus (including nucleus ventralis intermedius and nucleus ventralis oralis posterior) at St. George’s University Hospitals NHS Foundation Trust in London. All patients gave written informed consent, and the study was approved by the Health Research Authority UK and National Research Ethics Service (IRAS: 46576). DBS electrodes (Vercise Standard Lead, Boston Scientific) with eight ring contacts were implanted such that inferior contacts targeted STN, superior contacts targeted thalamus, and middle contacts spanned the rostral zona incerta. Data were collected 4–7 days postoperatively with externalized leads, while patients were off anti-dystonic medication. Continuous monopolar stimulation was delivered to middle contacts (C2–C7) at 130 Hz with linearly increasing amplitudes (0.5 to 4.5 mA or up to side effect threshold, in 0.5 mA steps), with each stimulation block lasting 46.92 *±* 0.99s (mean *±* SEM) and separated by 27.50 *±* 0.59s rest intervals.

Continuous EEG was recorded from six scalp channels (Cz, C3, C4, CPz, CP3, CP4) at 4096 Hz, and stimulus intensity was defined as the mean amplitude of resampled stimulation waveforms in non-overlapping 10-second bins. EEG data were band-pass filtered between 1 and 45Hz using a zero-phase FIR filter (8192 taps) to remove motion artifacts and high-frequency noise. Motion-contaminated segments were identified through a combination of fast step-like jumps (based on first differences exceeding three standard deviations) and slow baseline drifts (identified using a 2-second moving median and a similar threshold). Contaminated regions were padded by ±250ms, and only segments at least 100ms in duration were removed. The resulting clean EEG was downsampled to 500 Hz, epoched into 10-second trials, notch-filtered at 50, 100, 150, and 200Hz, and linearly detrended and demeaned. Trials and channels with high variance were excluded (*> µ* + *σ* for trials; *> µ* + 2.5*σ* for channels). Data were then re-referenced to the average, and independent component analysis was used to identify and remove up to three ocular components and one cardiac component per subject based on frequency power characteristics. A final trial-level exclusion was applied at *µ ±* 2*σ* to remove outlier trials. For each remaining segment, a deep neural network–based prediction of cortical state was computed. Stimulus amplitudes were matched to the corresponding EEG segments for all downstream analyses.

### Proximity to Edge-of-Chaos Criticality

To assess the chaoticity of the mean-field model’s dynamics (and whether its dynamics were near-critical), we calculated the stochastic largest Lyapunov exponent from simulated cortical LFPs. Lyapunov exponents measure how quickly nearby points in a system’s phase space diverge: a positive exponent indicates chaos (exponential divergence), a negative exponent suggests periodic or stable behavior (exponential convergence), and a near-zero exponent defines edge-of-chaos criticality. For each parameter set of our model, the stochastic Lyapunov exponent was estimated by running the model for 20 seconds with random initial conditions. At 9.999 seconds, a small random perturbation was applied, and the rate of divergence between the first run and the perturbed run was measured over the final 10 seconds. The divergence, *ϵ*(*t*), between the two runs was calculated as the squared difference:

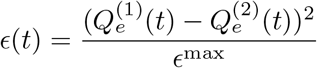

where *ϵ*^max^ is the maximum possible difference between the two runs:

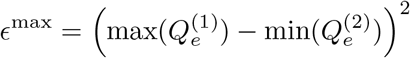

The largest Lyapunov exponent, Λ, was determined by fitting the exponential divergence *ϵ*(*t*):

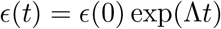

where *ϵ*(0) is the initial difference between 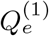 and 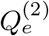 at *t* = 0. The slope of ln *ϵ*(*t*) versus *t* gives the estimate of the largest Lyapunov exponent. For all parameter configurations, both runs used identical noise inputs, meaning the slope of ln *ϵ*(*t*) versus *t* provides the stochastic Lyapunov exponent. Distance from edge-of-chaos criticality was then quantified as the absolute value of the stochastic largest Lyapunov exponent.

### Calculating spectral information transfer

Following our recent paper [1], we used the spectral information transfer measure developed by Pinzuti et al. [33], based on transfer entropy, which quantifies information transferred from a source variable *X* to a target variable *Y* [34–36]:

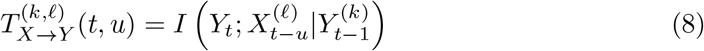

Here, *Y*_*t*_ is the state of *Y* at time 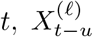 represents the past *ℓ* states of *X* with interaction delay *u*, and 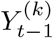 represents the past *k* states of *Y*. This measures how much uncertainty about *Y*_*t*_ is reduced by knowing 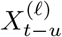, given 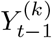. We used the Java Information Dynamics Toolkit (JIDT) [37] and, as described in our prior paper [1], we used the Kraskov method for probability distribution estimation using nearest-neighbor counting. As in our prior work [1], a fixed *K* = 4 nearest neighbors and *k* = *ℓ* = 1 were used in all datasets, and we scanned interaction delays *u* from 0.002 ms to 40 ms and selected the value that maximized transfer entropy.

Pinzuti et al.’s innovation allows estimation of information transfer in specific frequency bands using the maximum overlap discrete wavelet transform. Surrogate data are created by randomizing dynamics in specific frequency ranges (via Iterative Amplitude Adjustment Fourier Transform), enabling the estimation of spectral transfer entropy strength and statistical significance. The ‘swap-out swap-out’ (SOSO) algorithm, also from Pinzuti et al., identifies specific frequency bands for sending and receiving information between channels. We used SOSO for all spectral analyses and resampled data to 416 Hz for better alignment with canonical neural oscillations. Each analysis used 100 surrogates to ensure statistical significance and reliable spectral transfer entropy estimates.

### Entropy and complexity measures

Given the strong empirical association between level of consciousness and the entropy-/complexity of large-scale cortical electrodynamics [5, 38, 39], we likewise tested this relationship in our simulated model of DOC.

Permutation entropy (PE) was calculated to quantify the temporal complexity and irregularity in our time series data [40]. PE measures the entropy of ordinal patterns in a time series by comparing the order of neighboring values. For a time series 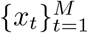, we constructed ordinal patterns of order *m* = 3 with time delays *τ ∈* {8, 16, 64}. For each embedding dimension *m* and time delay *τ*, we constructed vectors *X*_*t*_ = (*x*_*t*_, *x*_*t*+*τ*_, *x*_*t*+2*τ*_) for *t* = 1, 2, …, *M*− (*m* − 1)*τ*. Each vector *X*_*t*_ was mapped to one of the *m*! possible permutations *π* based on the relative order of its elements. The permutation entropy was then calculated as:

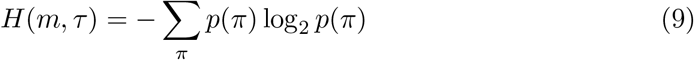

where *p*(*π*) represents the relative frequency of permutation pattern *π* in the time series. For equal values in the time series, we preserved the order of appearance when assigning ranks. The normalization of PE was performed by dividing by log_2_(*m*!), resulting in values between 0 and 1, where values close to 0 indicate highly regular signals and values close to 1 indicate highly complex or random signals. Different values of *τ* (8, 16, and 32, roughly corresponding to high-gamma, low-gamma, and beta waves, respectively) were employed to capture the entropy of dynamics occurring at different time scales.

Multiscale sample entropy (MSE) was also calculated to analyze the complexity of our time series data across multiple temporal scales [41]. MSE extends traditional sample entropy by incorporating a coarse-graining procedure to examine structural richness at different time scales. For our analysis, we set the embedding dimension *m* = 2, tolerance *r* = 0.15 times the standard deviation of the time series, and initial time scale *τ* = 1.

The MSE algorithm first constructs a coarse-grained time series 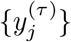 by averaging consecutive data points within non-overlapping windows of length *τ* :

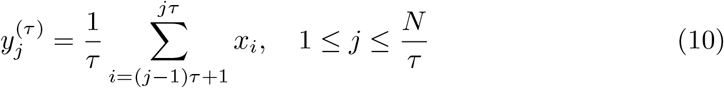

where {*x*_*i*_} represents the original time series and *N* is its length. For each coarse-grained time series, the sample entropy was calculated as:

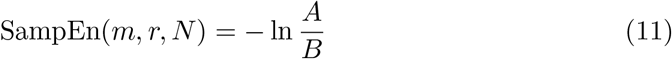

where *A* is the number of template vector pairs having a Chebyshev distance 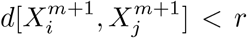 for vectors of length *m* + 1, and *B* is the number of template vector pairs having 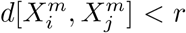 for vectors of length *m*. A lower value of sample entropy indicates more self-similarity in the time series, while higher values suggest greater irregularity and complexity.

Finally, Lempel-Ziv complexity (LZC) was also employed to quantify the randomness and complexity of our time series data [42]. LZC measures the rate of generation of new patterns in a sequence, with higher values indicating greater complexity.

The calculation of LZC first involved preprocessing the time series by detrending and demeaning the data, followed by binarization. Specifically, the instantaneous amplitude of the signal was computed using the Hilbert transform, and values greater than the mean amplitude were assigned 1, while the rest were assigned 0:

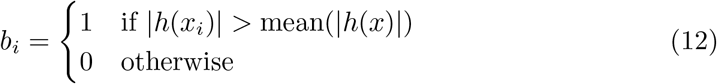

where *h*(*x*) represents the Hilbert transform of the original time series *x*. The resulting binary sequence was then analyzed using the Lempel-Ziv algorithm, which counts the number of distinct subsequences in the string when scanned from left to right. For a binary sequence *S* = *s*_1_*s*_2_ … *s*_*n*_, the algorithm decomposes the sequence into *c*(*n*) distinct patterns.

We normalized the LZC value by the asymptotic behavior of the complexity for a random sequence:

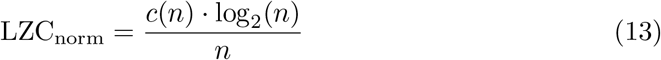

where *n* is the length of the time series. This normalization results in values between 0 and 1, with values approaching 1 indicating sequences with high complexity similar to random sequences, and values approaching 0 indicating highly regular sequences with patterns that repeat frequently.

