## Supplementary for "AI Discovery of Mechanisms of Consciousness, Its Disorders, and Their Treatment"

Supplementary Table 1: **Breakdown of training, validation, and test datasets used for training and evaluating our AI-based consciousness detectors.** Some subjects/animals contributed both cortical and subcortical recordings. African green monkeys contributed both training and validation data for the waking state, and independent test data for (induced) coma. ET=essential tremor, PD=Parkinson's disease, GPe=external globus pallidus, DOC=disorders of consciousness.

| **Dataset** | **Subjects** | **Samples (Train/** | **Val/** | **Test)** |
|---|---|---|---|---|
| **Subcortical Detectors Data (Training and Validation)** | | | | |
| Human ET Patient Thalamus | 10 | 470 | 118 | – |
| Human PD Patient GPe | 12 | 95 | 24 | – |
| Monkey GPe | 13 | 13,107 | 3,277 | – |
| Long-Evans Rat Thalamus | 9 | 200 | 50 | – |
| **Cortical Detector Data (Training and Validation)** | | | | |
| Human Acute Coma (TBI) | 22 | 602,696 | 150,674 | – |
| Human Chronic DOC | 14 | 49,062 | 12,266 | – |
| Human Healthy Control (dataset 1) | 10 | 7,066 | 1,766 | – |
| Human ET Patients | 10 | 354 | – | – |
| Human PD Patients | 12 | 51 | – | – |
| African Green Monkey (waking) | 13 | 7,656 | 1,914 | – |
| **Cortical Detector Data (Independent Test)** | | | | |
| Human Acute Coma (cardiac arrest) | 455 | – | – | 1,180,257 |
| Long-Evans Rat (waking and coma) | 9 | – | – | 250 |
| African Green Monkey (induced coma) | 13 | – | – | 9,570 |
| Human Healthy Control (dataset 2) | 20 | – | – | 3,674 |

Supplementary Table 2: **Firing-rate-related parameters for each population in our mean-field model of the conscious brain.** Parameters for the sigmoidal mean firing-rate response function include the membrane potential $\theta$ (mV) at which the population-average firing rate reaches half its maximum, the standard deviation $\sigma$ of the soma voltage relative to that threshold, and the maximum possible firing rate $Q_{\max}$ for that population.

| **Neural population** | Firing threshold $\theta$ (mV) | Voltage spread $\sigma$ (mV) | $Q_{\max}$ (spikes/s) |
|---|---|---|---|
| Excitatory | 6.03 | 2.9893 | 22.8294 |
| Inhibitory | 5.7626 | 6.1769 | 99.6704 |
| TRN | 0.59576 | 4.7428 | 385.3459 |
| Thalamic projection | 1.1724 | 9.6263 | 105.0741 |
| D1 | 1.1907 | 2.1266 | 19.3902 |
| D2 | 1.3843 | 4.5537 | 10.4114 |
| GPi | 3.53 | 22.244 | 132.0234 |
| GPe | 9.0081 | 2.2973 | 316.0468 |
| STN | 6.3887 | 3.406 | 202.4802 |

Supplementary Table 3: **Dendritic parameters for each connection in our mean-field model of the conscious brain.** The parameters $\alpha$ and $\beta$ are the decay and rise rates in 1/s, while $h_{\mathrm{syn}}$ is a synaptic scaling factor.

| **Connection** | $\alpha$ | $\beta$ | $h_{\mathrm{syn}}$ |
|---|---|---|---|
| Excitatory Cortical $\rightarrow$ Excitatory Cortical | 2.674 | 6.9705 | 19.555 |
| Inhibitory Cortical $\rightarrow$ Excitatory Cortical | 76.8488 | 91.4069 | 22.7117 |
| Thalamic Projection $\rightarrow$ Excitatory Cortical | 308.4148 | 309.811 | 14.0682 |
| Excitatory Cortical $\rightarrow$ Inhibitory Cortical | 23.8422 | 353.0609 | 45.2787 |
| Inhibitory Cortical $\rightarrow$ Inhibitory Cortical | 66.6434 | 185.5455 | 22.3397 |
| Thalamic Projection $\rightarrow$ Inhibitory Cortical | 53.1076 | 52.2119 | 5.4498 |
| GPe $\rightarrow$ Inhibitory Cortical | 49.1921 | 153.4782 | 44.3374 |
| Excitatory Cortical $\rightarrow$ TRN | 97.3268 | 258.7483 | 54.2651 |
| Thalamic Projection $\rightarrow$ TRN | 35.5804 | 1135.0763 | 15.9379 |
| GPe $\rightarrow$ TRN | 370.4681 | 420.044 | 11.7307 |
| Excitatory Cortical $\rightarrow$ Thalamic Projection | 82.407 | 185.5051 | 71.3342 |
| TRN $\rightarrow$ Thalamic Projection | 31.1955 | 217.8414 | 18.6017 |
| GPi $\rightarrow$ Thalamic Projection | 57.5824 | 89.063 | 25.6852 |
| GPe $\rightarrow$ Thalamic Projection | 136.9442 | 112.7362 | 8.971 |
| Excitatory Cortical $\rightarrow$ D1 | 393.5826 | 207.6264 | 32.657 |
| Thalamic Projection $\rightarrow$ D1 | 161.3917 | 64.0069 | 21.7262 |
| D1 $\rightarrow$ D1 | 118.2546 | 725.1904 | 20.9243 |
| GPe $\rightarrow$ D1 | 44.3807 | 134.2933 | 34.7326 |
| Excitatory Cortical $\rightarrow$ D2 | 66.4032 | 1787.148 | 82.5032 |
| Thalamic Projection $\rightarrow$ D2 | 81.9949 | 296.46 | 22.4305 |
| D2 $\rightarrow$ D2 | 30.8653 | 551.4931 | 14.9752 |
| GPe $\rightarrow$ D2 | 57.6319 | 359.2884 | 22.9981 |
| D1 $\rightarrow$ GPi | 2.544 | 341.8611 | 8.8214 |
| GPe $\rightarrow$ GPi | 53.9211 | 455.9377 | 19.0529 |
| STN $\rightarrow$ GPi | 95.4126 | 147.5733 | 14.2682 |
| Excitatory Cortical $\rightarrow$ GPe | 19.4579 | 322.3915 | 81.6792 |
| D2 $\rightarrow$ GPe | 4.8914 | 2382.7459 | 35.0419 |
| GPe $\rightarrow$ GPe | 1.5206 | 273.0801 | 44.268 |
| STN $\rightarrow$ GPe | 490.3135 | 1953.3756 | 22.4438 |
| Excitatory Cortical $\rightarrow$ STN | 33.1858 | 527.6786 | 16.9322 |
| GPe $\rightarrow$ STN | 189.6946 | 375.522 | 9.1586 |

Supplementary Table 4: **Propagator parameters for neural connections with coupling strengths.** Propagators can be *wave* or *map*. Wave-type propagators include a time-delay $\tau$, axonal spatial range $r$, and a damping coefficient $\gamma$. The parameter $\nu$ represents the axonal-synaptic coupling strength.

| **Connection** | **Type** | $\tau$ | $r$ | $\gamma$ | $\nu$ |
|---|---|---|---|---|---|
| Excitatory Cortical → Excitatory Cortical | Wave | 0.0037273 | 0.035009 | 52.7328 | 0.0001431 |
| Inhibitory Cortical → Excitatory Cortical | Map | 0.033308 | — | — | -0.0026139 |
| Thalamic Projection → Excitatory Cortical | Map | 0.042917 | — | — | 0.0024834 |
| Excitatory Cortical → Inhibitory Cortical | Wave | 4.8695e-5 | 0.084967 | 226.9079 | 0.0014598 |
| Inhibitory Cortical → Inhibitory Cortical | Map | 0.02991 | — | — | -0.0011039 |
| Thalamic Projection → Inhibitory Cortical | Map | 0.10776 | — | — | 0.00010874 |
| GPe → Inhibitory Cortical | Map | 0.016383 | — | — | -1.29e-05 |
| Excitatory Cortical → TRN | Wave | 0.01442 | 0.25159 | 29.0848 | 0.00027184 |
| Thalamic Projection → TRN | Map | 0.0044841 | — | — | 1.742e-05 |
| GPe → TRN | Map | 0.029018 | — | — | -0.00037503 |
| Excitatory Cortical → Thalamic Projection | Wave | 0.0063266 | 0.40024 | 55.2981 | 4.9136e-06 |
| TRN → Thalamic Projection | Map | 0.0030565 | — | — | -0.00069802 |
| GPi → Thalamic Projection | Map | 0.00072682 | — | — | -3.2149e-05 |
| GPe → Thalamic Projection | Map | 0.018695 | — | — | -0.00011238 |
| Stimulus → Thalamic Projection | Map | 0 | — | — | 0.0010994 |
| Excitatory Cortical → D1 | Wave | 0.00010577 | 0.025587 | 12.1336 | 0.00049354 |
| Thalamic Projection → D1 | Map | 0.003893 | — | — | 5.8137e-05 |
| D1 → D1 | Map | 0.0029817 | — | — | -0.00022084 |
| GPe → D1 | Map | 0.34971 | — | — | -6.4251e-05 |
| Excitatory Cortical → D2 | Wave | 0.0024002 | 0.059943 | 143.3842 | 0.00057826 |
| Thalamic Projection → D2 | Map | 0.00049981 | — | — | 2.1702e-06 |
| D2 → D2 | Map | 0.014893 | — | — | -0.00012266 |
| GPe → D2 | Map | 0.0099476 | — | — | -0.00018428 |
| D1 → GPi | Map | 0.00056918 | — | — | -3.0066e-05 |
| GPe → GPi | Map | 0.00072232 | — | — | -4.2334e-06 |
| STN → GPi | Map | 0.0017869 | — | — | 1.7415e-05 |
| Excitatory Cortical → GPe | Wave | 0.016993 | 0.041773 | 95.3548 | 5.7408e-05 |
| D2 → GPe | Map | 0.00014521 | — | — | -0.00024693 |
| GPe → GPe | Map | 0.003942 | — | — | -7.3175e-05 |
| STN → GPe | Map | 0.00061743 | — | — | 0.00036479 |
| Excitatory Cortical → STN | Wave | 0.0016426 | 0.38859 | 223.5338 | 0.00025665 |
| GPe → STN | Map | 0.00047147 | — | — | -5.9005e-05 |

Supplementary Table 5: **Mean firing rates (in spikes/s) of each brain region in the mean-field simulation of the conscious brain, compared to empirical ranges of firing rates from multiple mammalian species.** The firing rates for each simulated brain region were optimized, along with the outputs of the consciousness-detector networks, through a genetic algorithm to align with known physiological ranges. The GPi and SNr were treated as a single population, sharing the same firing rate. GPi=internal globus pallidus, SNr=substantia nigra pars reticulata, GPe=external globus pallidus, STN=subthalamic nucleus, TRN=thalamic reticular nucleus.

| | Simulation of Conscious Brain | Monkeys | Rats | Mice | Humans | Cats |
|---|---|---|---|---|---|---|
| Cortical Pyramidal Cells | 6.99 | 5-20[a] | 2-5[b] | 2-11[c] | 1-10[d] | |
| Cortical Inhibitory Interneurons | 18.21 | 0.1-100[e] | | 1-50[f] | | |
| Striatum | 4.82 | 4-7[g] | 1-7[h] | | | |
| GPi | 60.77 | 60-90[i] | 15-20[j] | | | |
| SNr | 60.77 | 50-70[k] | | | | |
| GPe | 43.74 | 16-70[l] | 16-115[m] | 1-64[n] | | |
| STN | 22.16 | 20-30[o] | 8-11[p] | | | |
| Projection Nuclei | 14.10 | 5-25[q] | | | 10-20[r] | |
| TRN | 24.10 | | | 4-64[s] | | 20-30[t] |

a 1,2

b 3

c 4

d 5

e 6

f 7

g 1,8

h 3,9

i 10–13

j 3

k 14,15

l 10,11,13,16,17

m 18,19

n 20

o 11,21

p 22

q 23

r 24

s 25–28

t 29

Supplementary Table 6: **Pairwise discrimination of subcortical LFPs via relative band-power and zero-crossing.** For each region-pair, we applied one-tailed unequal-variance $t$-tests to the z-scored real LFPs (from bats & GAERS rats for striatum; Long–Evans rats & ET patients for thalamus; African green monkeys & PD patients for GPe) and to the corresponding simulated LFPs (GPe/striatum/thalamus only, 600 simulated 10-second LFP samples per region). We tested whether region A $>$ region B on feature $f$ if the real-data mean difference $\Delta_{\text{Real}} = \mu_A - \mu_B$ was positive (otherwise A<B). All $p$-values are Bonferroni-corrected over 12 tests ($3$ pairs×$4$ features). Stars denote $p < 0.05$. These results show that, with the exception of delta power, the real-world features that distinguish LFPs from different subcortical regions are recapitulated in the model—even though we never explicitly programmed them—paralleling our successful modeling of region-specific firing rates (Table S5).

| Pair | Feature | $\Delta_{\text{Real}}$ | $\Delta_{\text{Sim}}$ | $p^{\text{BF}}_{\text{Real}}$ * | $p^{\text{BF}}_{\text{Sim}}$ * |
|---|---|---|---|---|---|
| Striatum vs GPe | $\delta_{\text{rel}}$ | $-0.042$ | $0.062$ | $4.70 \times 10^{-4}$* | $1.00$ |
| | $\theta_{\text{rel}}$ | $-0.067$ | $-0.062$ | $1.05 \times 10^{-15}$* | $3.69 \times 10^{-82}$* |
| | $\alpha_{\text{rel}}$ | $-0.109$ | $-0.0001$ | $1.00 \times 10^{-37}$* | $1.00$ |
| | zcr | $-0.011$ | $-0.006$ | $6.24 \times 10^{-21}$* | $5.62 \times 10^{-205}$* |
| Striatum vs Thal | $\delta_{\text{rel}}$ | $0.093$ | $-0.012$ | $1.04 \times 10^{-17}$* | $1.00$ |
| | $\theta_{\text{rel}}$ | $0.054$ | $0.033$ | $1.40 \times 10^{-11}$* | $7.10 \times 10^{-26}$* |
| | $\alpha_{\text{rel}}$ | $-0.147$ | $-0.021$ | $3.49 \times 10^{-59}$* | $2.46 \times 10^{-17}$* |
| | zcr | $-0.013$ | $-0.042$ | $5.61 \times 10^{-28}$* | 0.00* |
| GPe vs Thal | $\delta_{\text{rel}}$ | $0.135$ | $-0.075$ | $7.37 \times 10^{-229}$* | $1.00$ |
| | $\theta_{\text{rel}}$ | $0.121$ | $0.095$ | $1.27 \times 10^{-262}$* | $1.27 \times 10^{-158}$* |
| | $\alpha_{\text{rel}}$ | $-0.257$ | $-0.021$ | $0$* | $1.49 \times 10^{-16}$* |
| | zcr | $-0.0016$ | $-0.037$ | $1.13 \times 10^{-2}$ | 0.00* |

Supplementary Table 7: **Additional model parameters significantly associated with AI-predicted level of consciousness in simulated DOC**. Model parameters were identified using ridge-penalized linear regression with inverse Gaussian transformation and permutation testing (1,000 permutations), followed by False Discovery Rate (FDR) correction. Each coefficient reflects the standardized association between a feature and predicted level of consciousness.

| **Additional Model Parameters Associated with AI-Predicted Level of Consciousness** | **Coefficient** | *p-value (Permutation test, FDR-corrected)* |
|---|---|---|
| Striatum (D2-expressing) to Striatum (D2-expressing) Propagation Delay | -0.172 | $<$0.0001 |
| Thalamic Projection Nucleus to TRN Propagation Delay | -0.171 | $<$0.0001 |
| Inhibitory to Inhibitory Cortical PSP Height | -0.161 | $<$0.0001 |
| Excitatory Cortex to Striatum (D2-expressing) Propagation Delay | -0.153 | $<$0.0001 |
| Thalamic Projection Nucleus Cross-Neuron Variance of Subthreshold Voltages | +0.133 | $<$0.0001 |
| GPe to Striatum (D2-expressing) PSP Height | +0.131 | $<$0.0001 |
| Excitatory Cortical to Striatum (D2-expressing) PSP Height | -0.126 | $<$0.0001 |
| Excitatory Cortical Threshold Potential | +0.117 | $<$0.0001 |
| TRN Threshold Potential | +0.114 | $<$0.0001 |
| Excitatory to Inhibitory Cortical Propagation Damping | +0.105 | $<$0.0001 |
| Striatum (D1-expressing) Variance of Subthreshold Voltages | -0.105 | 0.0058 |
| GPe to TRN PSP Height | +0.104 | 0.0100 |
| Inhibitory to Excitatory Cortical PSP Height | +0.103 | $<$0.0001 |
| GPe to Striatum (D1-expressing) Propagation Delay | +0.102 | 0.0058 |
| Thalamic Projection Nucleus Threshold Potential | -0.101 | 0.0180 |
| GPe to GPi PSP Height | +0.100 | 0.0200 |
| TRN to Thalamic Projection Nucleus PSP Duration | -0.096 | $<$0.0001 |
| STN to GPi PSP Duration | +0.094 | 0.0100 |
| Excitatory Cortical to GPe Propagation Damping | -0.094 | 0.0058 |
| GPe to Striatum (D1-expressing) PSP Height | -0.094 | 0.0230 |
| Excitatory to Inhibitory Cortical Propagation Range | -0.093 | 0.0180 |
| Excitatory Cortical to Striatum (D2-expressing) Cortical PSP Duration | +0.091 | 0.0200 |
| Striatum (D2-expressing) to GPe Propagation Delay | +0.116 | 0.0100 |
| STN to GPe PSP Height | -0.087 | 0.0180 |
| GPe to STN PSP Duration | -0.084 | 0.0180 |
| GPe to GPi Propagation Delay | -0.081 | 0.0200 |
| GPe to Thalamic Projection Nucleus Propagation Delay | -0.080 | 0.0280 |
| GPe to TRN Propagation Delay | +0.079 | 0.0460 |
| Excitatory to Inhibitory Cortical Propagation Delay | -0.071 | 0.0490 |

Supplementary Table 8: **Partitioning of explained variance (total $R^2 = 0.205$) for left striatum–GPe streamline counts.** Each predictor's value is the percentage of the total explained variance uniquely attributable to that factor in a linear model including diagnosis (VS vs. MCS/eMCS), age at MRI, gender, days post-injury, and injury etiology.

| **Predictor** | **% of Total $R^2$** |
|---|---|
| Diagnosis (VS vs. MCS/eMCS) | 35.0 |
| Age at MRI | 8.9 |
| Gender | 12.6 |
| Days post-injury | 5.8 |
| Etiology | 37.8 |
| **Total $R^2$** | 0.205 |

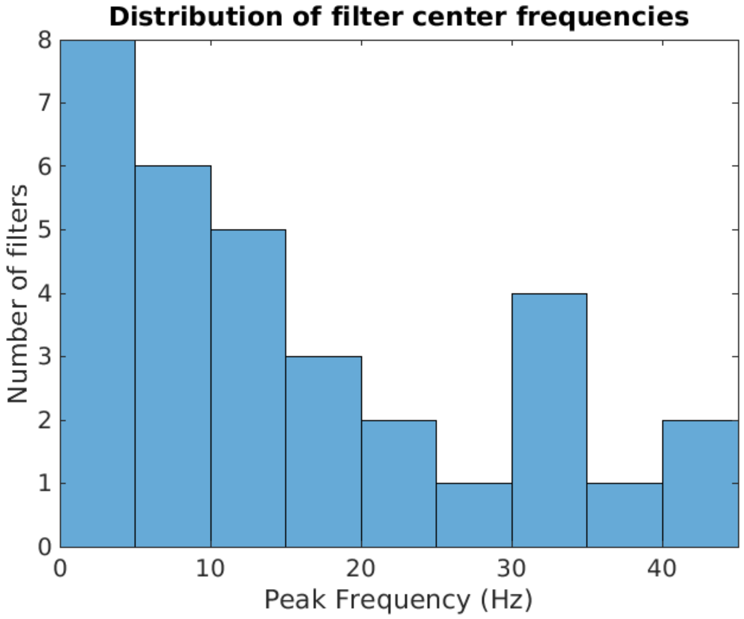


Supplementary Figure 1: **Distribution of learned temporal-kernel tuning in the cortical consciousness detector DCNN.** The histogram shows the peak-frequency tuning of the 32 one-dimensional kernels in the first convolutional layer of our deep convolutional neural network, trained to detect consciousness from cortical electrophysiology recordings. For each kernel, we computed its Fourier-domain gain (0.5–45 Hz) and identified the single frequency at which that gain was maximal, then binned those peak frequencies into 5 Hz intervals. The majority of kernels (19/32) tune to low-frequency oscillations (0–15 Hz), reflecting the known importance of slow and mid-range rhythms in disorders of consciousness. A smaller secondary cluster (4/32) peaks in the 30–35 Hz range, suggesting sensitivity to $\beta$ activity. Importantly, these kernels do not only measure spectral power—they act as matched filters, detecting the specific waveform shapes (e.g. burst-like envelopes and phase structure) within each band that best discriminate conscious from comatose states.

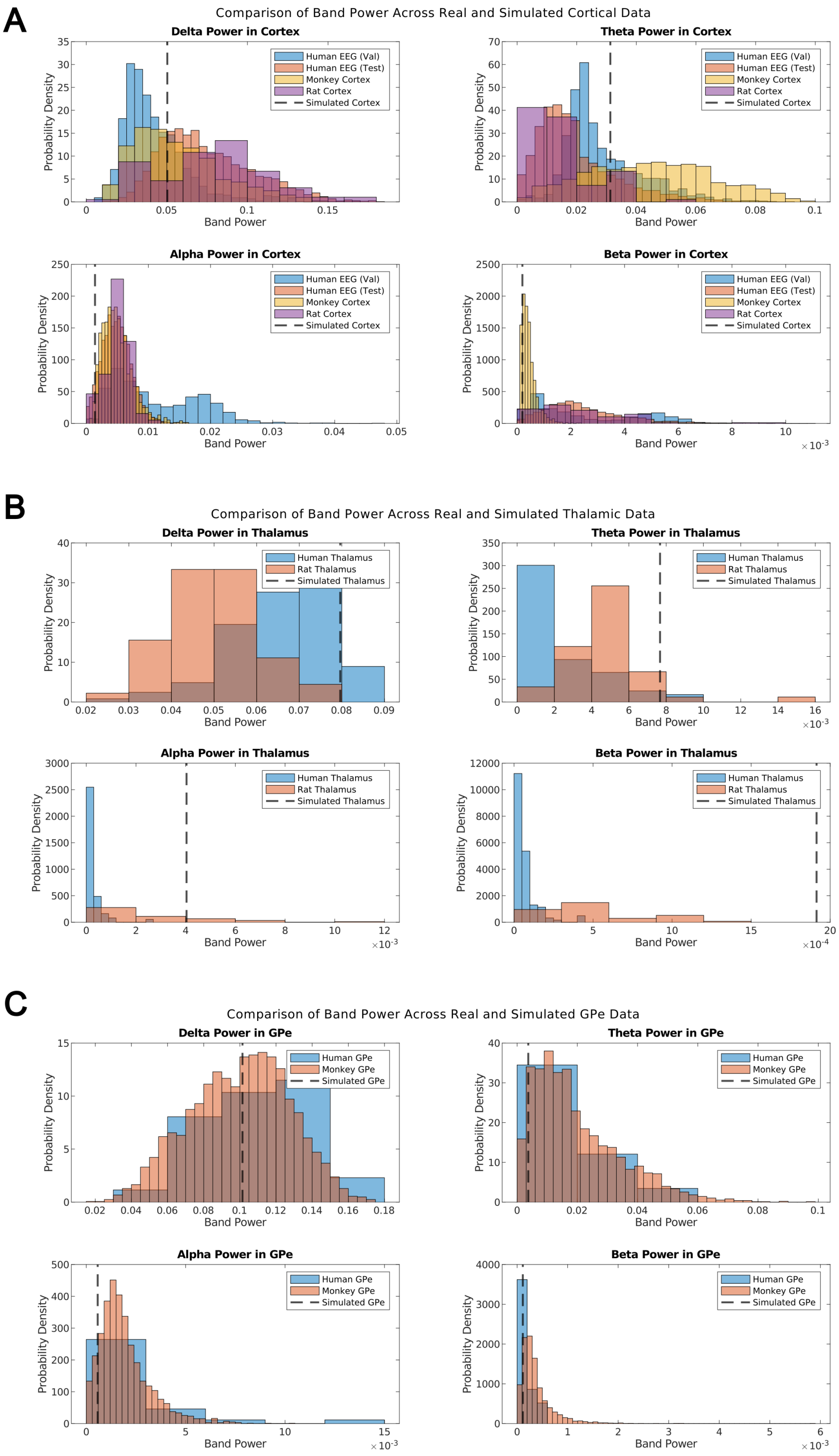


Supplementary Figure 2: **Comparison of band power across simulated and real LFP, ECoG, and EEG data from cortex, thalamus, and GPe.** To compare simulated to real data across species and modalities, we analyzed band-specific power in four canonical frequency bands: delta (0.5–3 Hz), theta (4–7 Hz), alpha (8–12 Hz), and beta (13–25 Hz). Power spectra for all z-scored time series were computed using Welch's method, and mean power was calculated across all frequency bins within each band. Histograms show the distribution of band power across all available held-out validation or independent test samples from real recordings for each region, with each color indicating a different dataset. Vertical dashed lines indicate the corresponding band power for the simulated region. Panel A shows results for cortex, panel B for thalamus, and panel C for GPe. These results show that, with the exception of beta power in the simulated thalamic LFP, the spectral properties of the simulated cortical, thalamic, and pallidal LFPs all fall within physiological ranges.

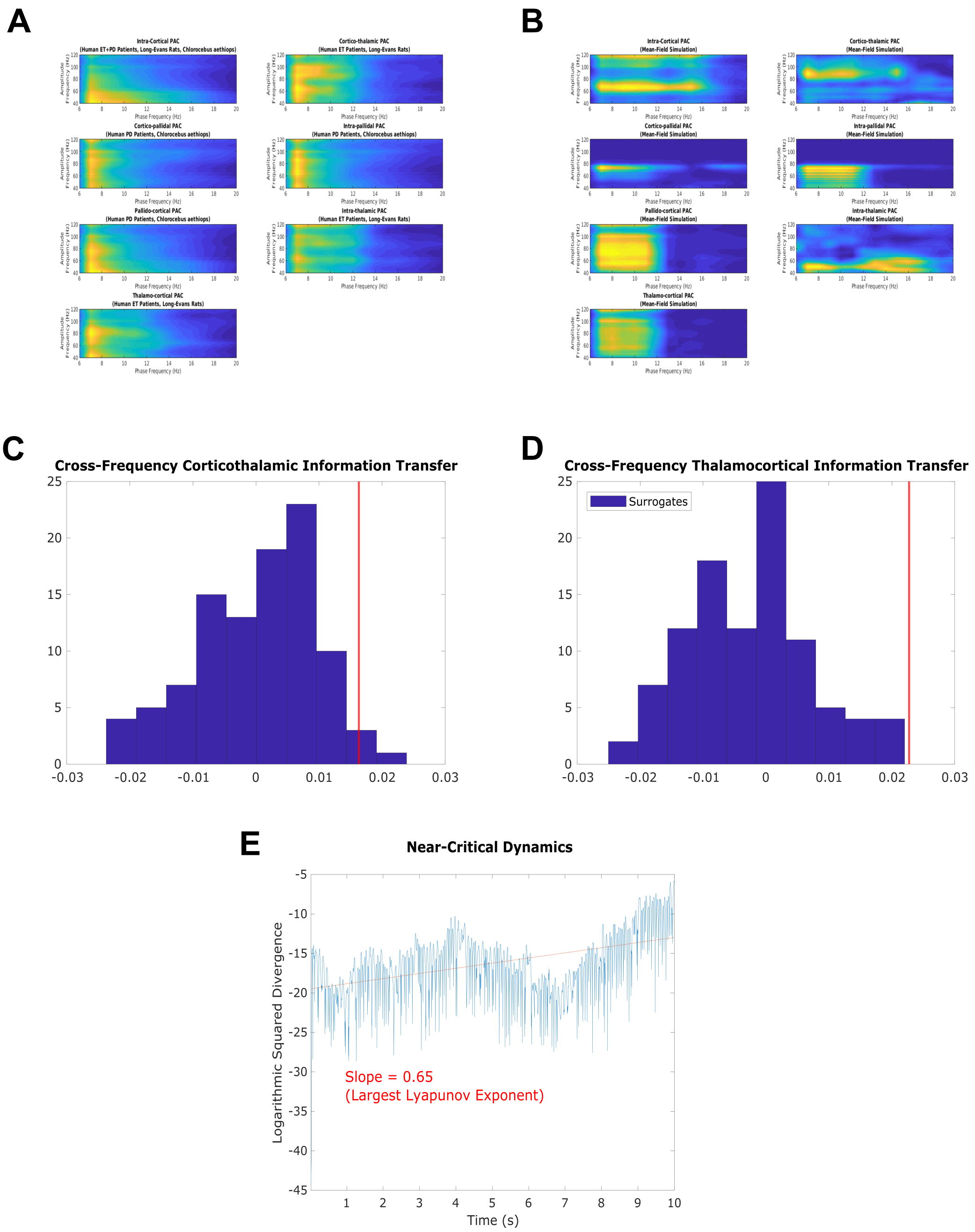


Supplementary Figure 3: **Simulated brain dynamics recapitulate key electrophysiological signatures of consciousness, including cross-frequency coupling, directed information flow, and near-critical chaotic dynamics.** **(A-B)** Phase-amplitude coupling in real (E) versus simulated (F) LFPs. In both real and simulated data, strong theta-gamma coupling was observed within the cortex, thalamus, and GPe, as well as between brain regions. However, simulated data exhibited stronger alpha-gamma coupling. **(C-D)** Directed information transfer from low-frequency (∼1–13 Hz) cortical rhythms to high-frequency (52–104 Hz) thalamic rhythms, and vice versa. The red vertical line represents the difference between the calculated transfer entropy from source to target when signals are unscrambled, and the estimated transfer entropy when low-frequency dynamics at the source are scrambled. The blue histogram represents the distribution of values across 100 surrogate datasets, where both low-frequency source and high-frequency target dynamics are scrambled. Statistical significance is determined if fewer than 5% of surrogate values (blue histogram) exceed this difference (red line) (see Methods for further details). **(E)** Weak chaos in the simulation of a cortical LFP in the conscious brain, indicated by the logarithmic squared divergence between two 10-second runs of the model with slightly different initial firing rates. The slope of the fitted line is positive but near-zero, indicating weakly chaotic simulated cortical dynamics near edge-of-chaos criticality.

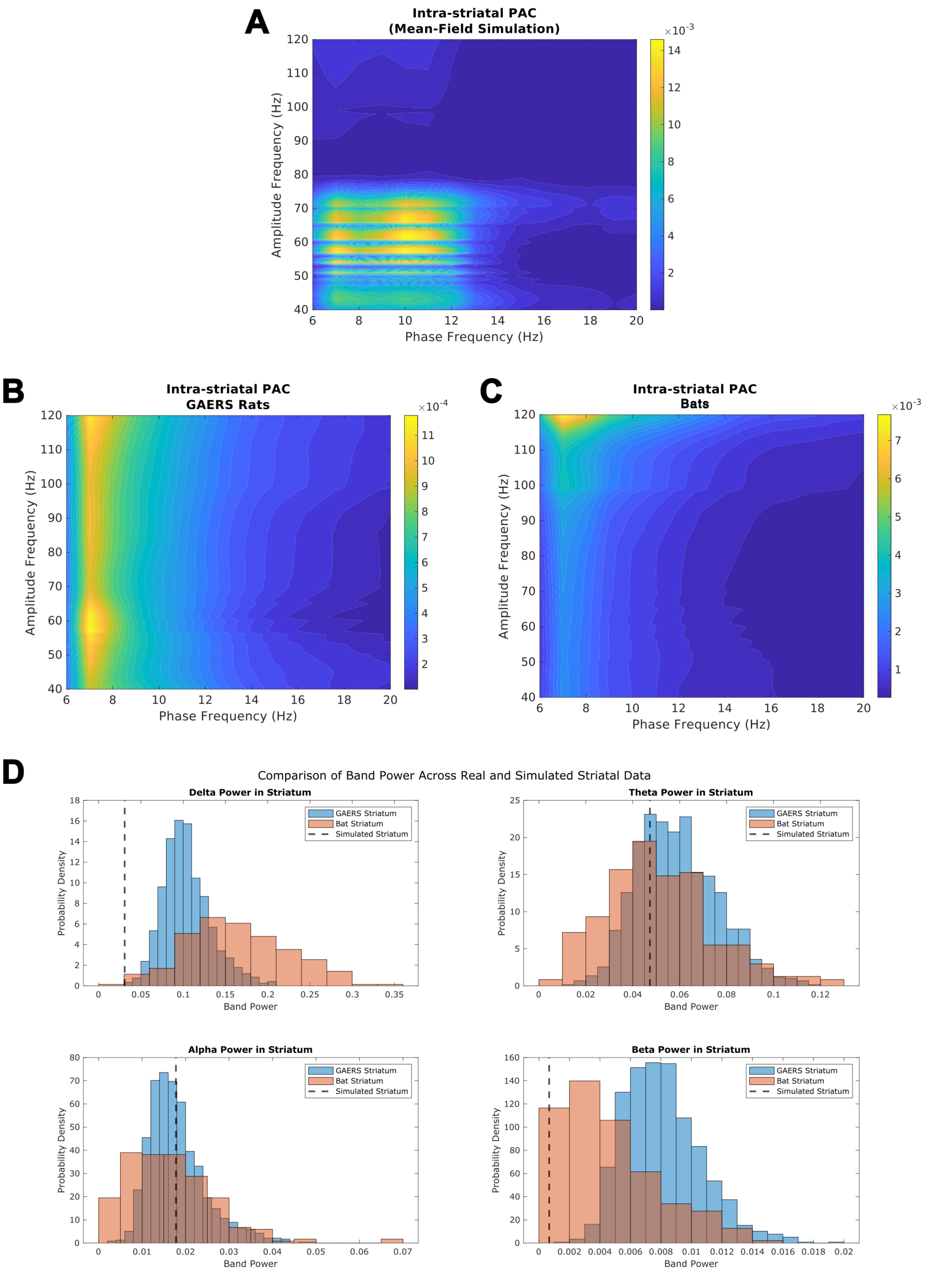


Supplementary Figure 4: **Validation of simulated striatal LFPs against real striatal recordings from multiple species.** Although our model was explicitly trained to reproduce electrophysiological features of the cortex, thalamus, and GPe, it was not trained on any striatal recordings. Here, we validate the realism of the model's simulated striatal LFP by comparing it to empirical recordings from the dorsal striatum of two species: Genetic Absence Epilepsy Rats from Strasbourg (GAERS) and Seba's short-tailed bats (*Carollia perspicillata*). **(A)** Phase–amplitude coupling (PAC) between low-frequency phase and high-frequency amplitude in the model's striatal LFP revealed robust theta–gamma coupling, along with alpha-gamma coupling. **(B–C)** Comparable intra-striatal PAC patterns in GAERS rats (B) and bats (C) show a similar concentration of theta–gamma coupling. **(D)** Band-specific power distributions in delta, theta, alpha, and beta frequency bands from the model's simulated striatum (vertical dashed lines) closely matched those observed in empirical striatal data from both GAERS rats (blue) and bats (orange), despite no model exposure to these datasets. Together, these results suggest that the model successfully generalizes to unseen brain regions, producing biologically realistic LFP dynamics in the striatum.

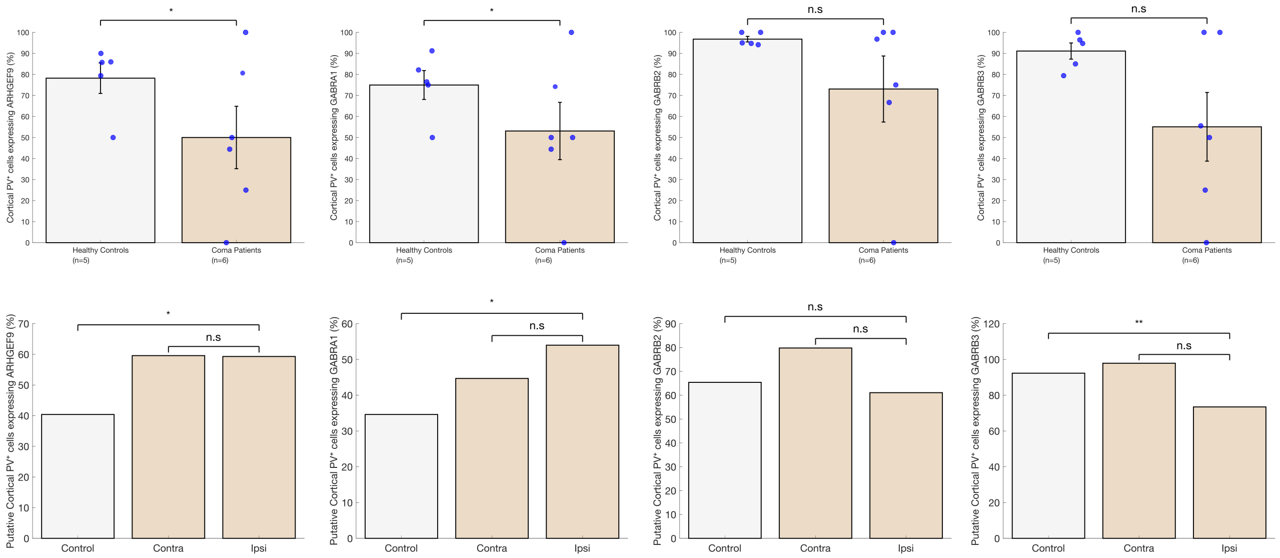


Supplementary Figure 5: **Expression of additional inhibitory synapse-related genes in cortical PV⁺ interneurons from human coma patients and a rat model of severe ischemic stroke. Top row:** Percentage of cortical PV⁺ cells expressing *ARHGEF9*, *GABRA1*, *GABRB2*, and *GABRB3* in non-neurological controls (light gray) and coma patients (tan) based on human snRNAseq data (n = 5 controls, n = 6 patients). Blue dots indicate individual donor values. Bars: mean ± SEM. One-tailed generalized linear mixed-effects models (GLMMs) were used to assess group differences, with donor as a random effect. Significant down-regulation in coma was observed for *ARHGEF9* and *GABRA1*; no significant change was observed for *GABRB2* or *GABRB3*. **Bottom row:** Percentage of putative cortical PV⁺ cells expressing the same four genes in sham controls (light gray), and in contralateral and ipsilateral hemispheres following severe ischemic stroke (tan) based on rat snRNAseq data (n = 1 sham, n = 1 paired stroke). Putative cortical PV⁺ cells were defined by co-expression of *Pvalb* and *Satb1*, excluding cells expressing subcortical markers *Pthlh* or *C1ql1*. Bars: mean. One-tailed GLMMs were used at the cell level to assess expression changes. Significant up-regulation was observed for *ARHGEF9*, *GABRA1*, and *GABRB3* in the hemisphere ipsilateral to the ischemic lesion compared to sham. In sum, no consistent pattern of regulation was observed for these genes across species and injury etiologies. *$p < 0.05$, **$p < 0.01$, n.s. not significant.

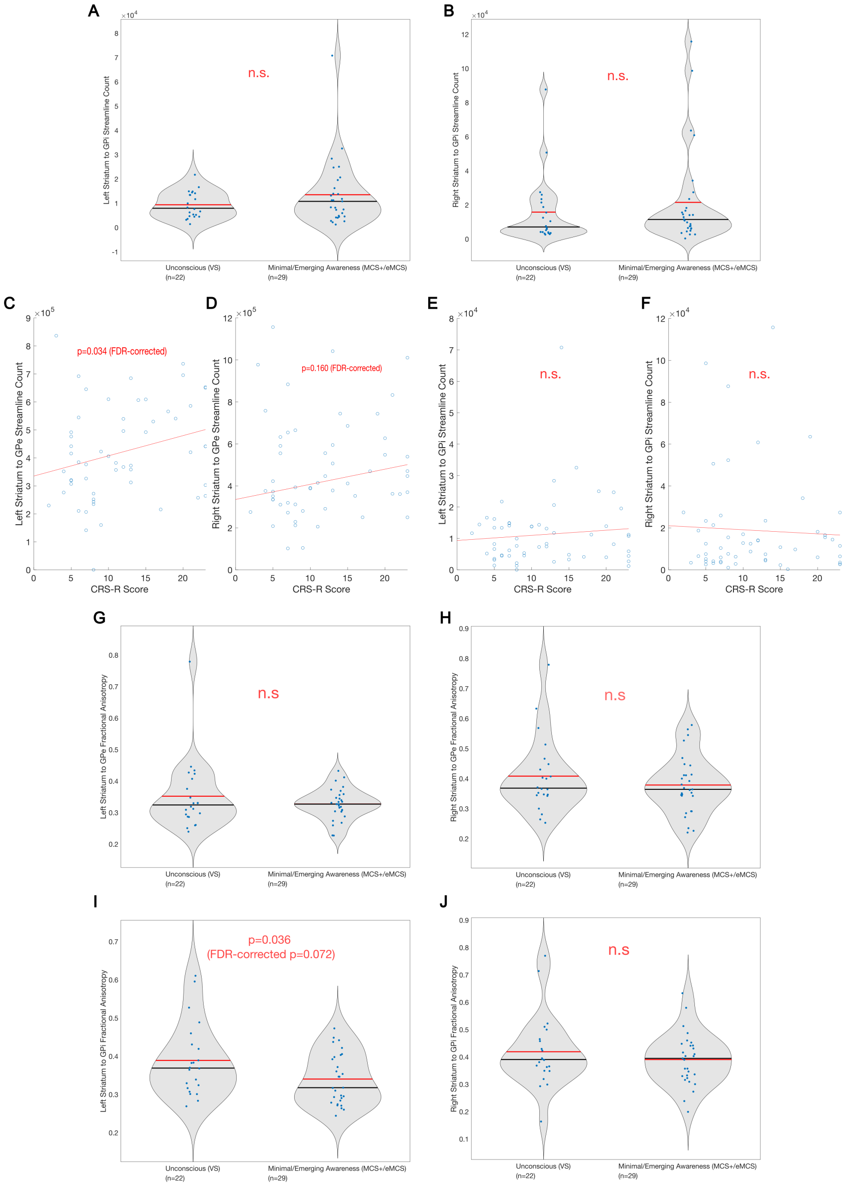


Supplementary Figure 6: **Supplemental analysis of striato-pallidal structural connectivity in disorders of consciousness.** Panels (A–B) show no significant group differences in striatum-to-GPi streamline counts between vegetative state (VS) and minimally conscious/emergent patients (MCS+/eMCS) in either the left (A) or right hemisphere (B) (Wilcoxon rank-sum test). Panels (C–D) show a significant positive Spearman correlation between left striatum–GPe streamline count and CRS-R score (C; $\rho = 0.263$, 1,000 permutation bootstrap $p = 0.034$, FDR-corrected for two comparisons across the left and right hemispheres), but no significant correlation in the right hemisphere (D). Panels (E–F) show no significant correlations between striatum–GPi streamline counts and CRS-R scores in either hemisphere. Panels (G–H) show no significant group differences in fractional anisotropy (FA) along striatum–GPe tracts. Panel (I) shows a significant increase in FA along the left striatum–GPi tract in VS patients compared to MCS+/eMCS patients ($p = 0.036$, FDR-corrected $p = 0.072$, one-tailed Wilcoxon rank-sum test), while panel (J) shows no significant group difference in FA along the right striatum–GPi tract. Together, these results support selective disruption of the striatum–GPe pathway in unconsciousness, with perhaps pathological reinforcement of left striatum–GPi tracts in VS, as predicted by our AI-driven model.

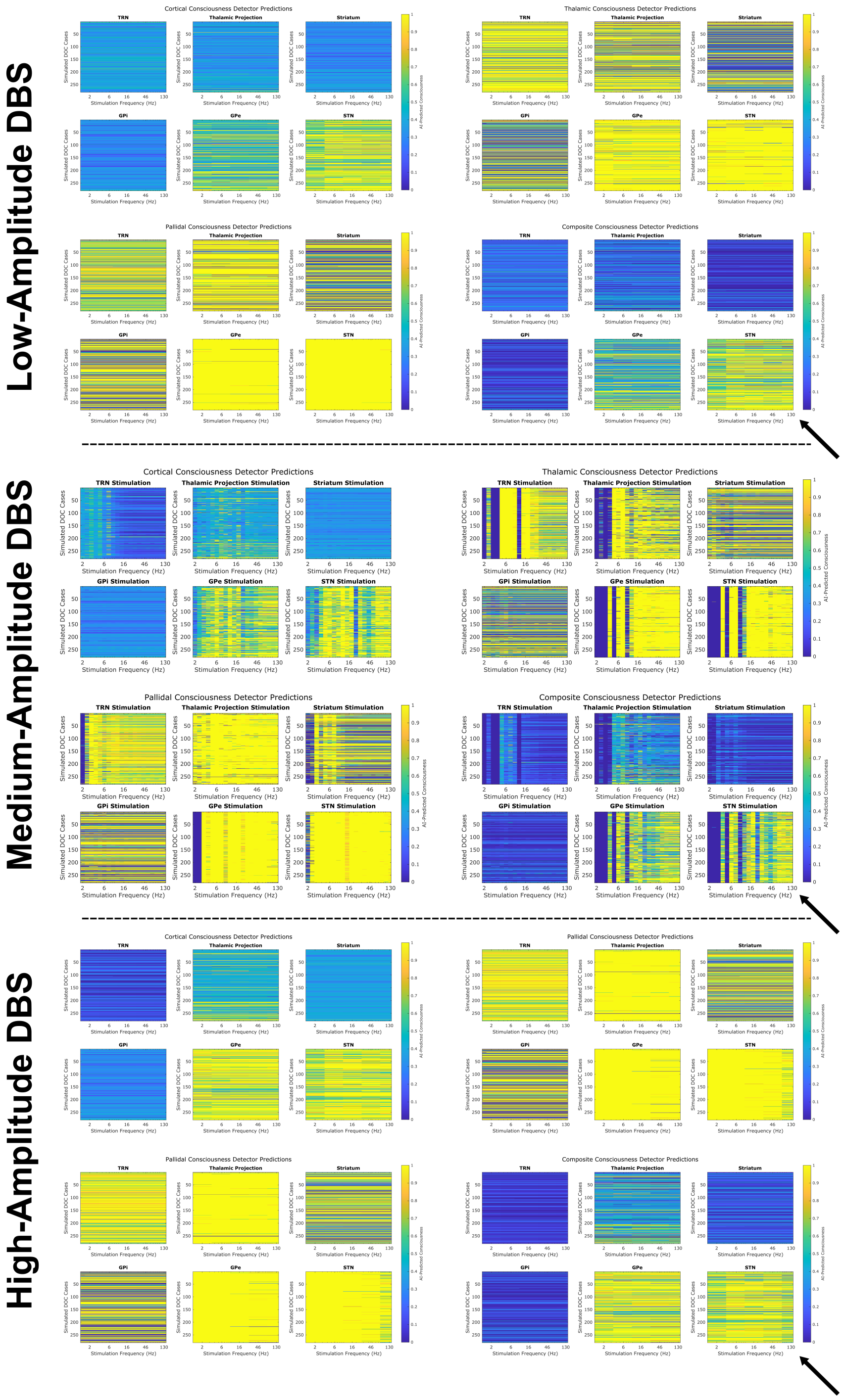


Supplementary Figure 7: **Effects of simulated DBS across subcortical targets, frequencies, and amplitudes on the predictions of our cortical, thalamic, pallidal, and composite consciousness detectors.** Here, low-amplitude simulated DBS corresponded to setting the amplitude of the stimulatory neural population to 5 (arbitrary units), medium-amplitude DBS corresponded to an amplitude of 10, and high-amplitude DBS corresponded to an amplitude of 15. Note that in Fig. 5 in the main paper, we report the results of medium-amplitude DBS. Importantly, high-frequency subthalamic stimulation emerged as the optimal predicted modality for restoring consciousness (estimated by our composite predictor) across different simulated DBS amplitudes (diagonal arrows).

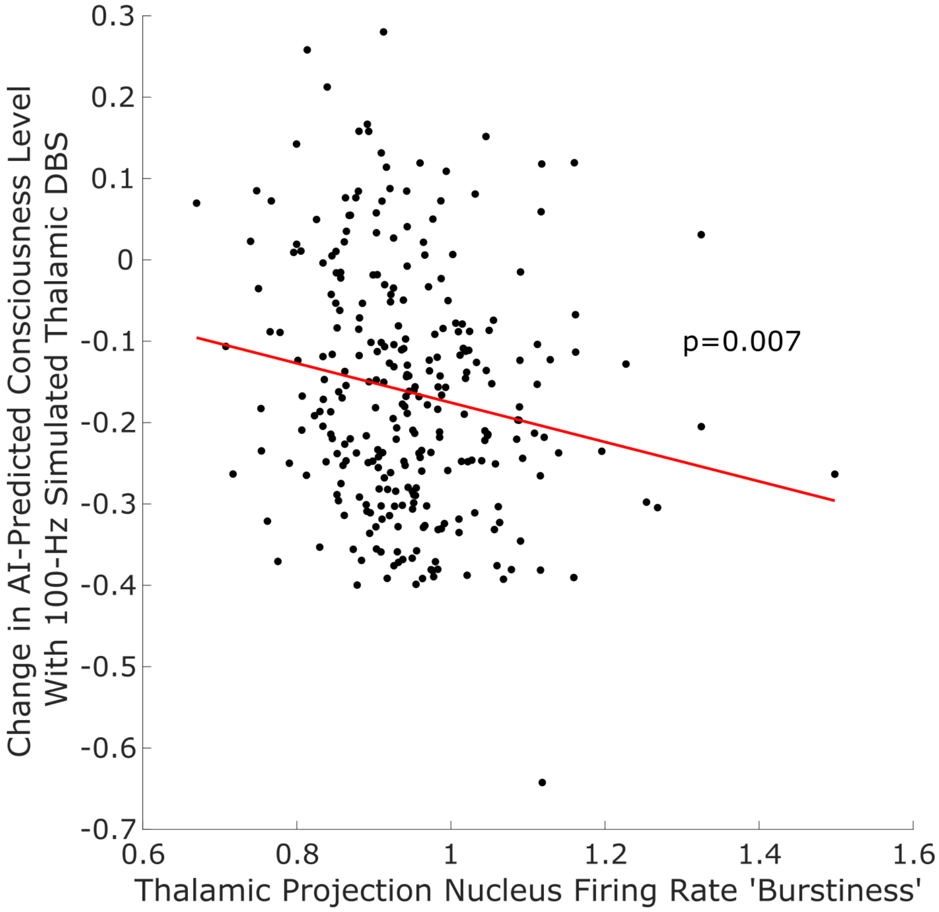


Supplementary Figure 8: **Relationship between thalamic burst-duration variability and change in AI-predicted level of consciousness (from our trained cortical consciousness detector) under 100 Hz simulated deep brain stimulation to the thalamic projection nucleus.** X-axis: Coefficient of variation of burst durations, $\mathrm{CV}_{\mathrm{burst}} = \frac{\sigma_T}{\mu_T}$, where burst durations $T$ are the lengths (in seconds) of contiguous epochs for which the mean thalamic firing rate exceeded $\mu_r + 2\,\sigma_r$ (with $\mu_r$ and $\sigma_r$ the mean and standard deviation of the entire rate trace). Y-axis: Change in the AI-predicted cortical consciousness level relative to baseline during 100 Hz stimulation. Each dot is one simulation trial; the red line is the least-squares fit (slope=-0.2419). The p-value (p = 0.007) was computed by a 1,000-iteration permutation test on the slope coefficient. The negative association between thalamic burstiness and "recovery" of consciousness succesfully retrodicts the recent finding that more tonic thalamic firing is predictive of responsiveness to 100 Hz intralaminar thalamic DBS in DOC.
